# SeroBA(v2.0) and SeroBAnk: a robust genome-based serotyping scheme and comprehensive atlas of capsular diversity in Streptococcus pneumoniae

**DOI:** 10.1101/2025.04.16.648953

**Authors:** Oliver Lorenz, Alannah C. King, Harry C. H. Hung, Feroze A. Ganaie, Anne L. Wyllie, Sam Manna, Catherine Satzke, Mark van der Linden, Neil Ravenscroft, Hans-Christian Slotved, Lesley McGee, Moon H. Nahm, Stephen D. Bentley, Stephanie W. Lo

## Abstract

The unprecedented number of *Streptococcus pneumoniae* (the pneumococcus) genomes sequenced in recent years has accelerated the discovery of novel serotypes and highlighted the genetic diversity both between and within each serotype. A novel serotype should demonstrate a distinct *cps* locus, capsular structure, and serological profile. In only the past four years, nine new serotypes have been identified. Accurate and timely serotyping of pneumococcal isolates is key to understanding its global distribution, evolution, and the response of the bacterial population to vaccination. However, current bioinformatics serotyping tools are infrequently updated, and struggle to accommodate the rapid discovery of new serotypes in a timely manner. To address these limitations, we built a comprehensive and curated library (SeroBAnk) encompassing all known pneumococcal serotypes; this resource is presented as an atlas on a dedicated publicly accessible webpage (https://www.pneumogen.net/gps/#/serobank). Building upon this resource, we developed SeroBA(v2.0), a tool with an easy-to-update database that can accurately identify 102 of 107 known pneumococcal serotypes (except for serotypes 24B, 24C, 24F, 7D and 6H) and 18 genetic subtypes within serotypes 6A, 6B, 11A, 19A, 19F and 33F. We validated SeroBA(v2.0) on 26,306 genomes from the Global Pneumococcal Sequencing project, reference isolates and simulated reads derived from the reference genetic sequences of capsular polysaccharide biosynthetic (*cps*) locus and showed that SeroBA(v2.0) can reliably detect the nine recently discovered serotypes. Additionally, we show that *in silico* serotypes inferred by SeroBA(v2.0) had high concordance with phenotypic serotypes determined by either Quellung or latex agglutination at the serotype level (88.9%; 15,945/17,933), and at the serogroup level (91.9%; 16,480/17,933). Finally, we propose a community-contribution based approach to ensure that SeroBA(v2.0) is maintained and updated as novel serotypes continue to be discovered. The global community can submit putative novel serotypes through our public repository on GitHub (https://github.com/GlobalPneumoSeq/seroba/issues). The submitted putative novel serotypes will be curated based on the genetic sequence of *cps* region, capsular structure and serological profile by people of relevant expertise in the field. SeroBA(v2.0) can be accessed at https://github.com/GlobalPneumoSeq/seroba.

**Data summary:** Genome sequences are available in the European Nucleotide Archive (ENA) and are also available alongside metadata on the Monocle Database available at https://data.monocle.sanger.ac.uk/. The authors confirm all supporting data, code and protocols have been provided within the article or through supplementary data files.

**Impact Statement:** The polysaccharide capsule has been an effective vaccine antigen against diseases caused by *Streptococcus pneumoniae* (the pneumococcus). The pneumococcal conjugate vaccine has been estimated to have halved pneumococcal-related childhood mortality over 15 years (2000-2015). We collated the genetic locus and capsular structure of each known capsule type (serotype), alongside with pneumococcal vaccine formulation and licensure history, into a single webpage (SeroBAnk), providing a valuable resource for basic research and vaccine development. With increasing use of whole genome sequencing in clinical and public health laboratories, we also provided a fast and accurate bioinformatics tool, SeroBA(v2.0), to identify 102 pneumococcal serotypes, alongside a proposed system to expand SeroBA(v2.0) to include new serotypes as they are discovered, ensuring that the tool remains valuable to the global research community in the long-term.

## Introduction

*Streptococcus pneumoniae* (the pneumococcus) is a gram-positive bacterial pathogen which can cause diseases such as pneumonia, sepsis and meningitis. Despite effective vaccines that have been estimated to have reduced pneumococcal deaths in children under five years of age by 51% between 2000 and 2015 [1], pneumococcus is still estimated to have caused ∼318,000 global deaths in 2015 [1]. Pneumococcus is typically surrounded by a polysaccharide capsule, which facilitates the evasion of opsonophagocytosis [2], which is one of the main reasons why the polysaccharide capsule is considered to be a major virulence factor [2].

The structure of the capsule is determined by the genes of the capsular polysaccharide biosynthetic locus (*cps*), which encodes for proteins such as transferases, polymerases, and flippases [3]. In general, pneumococcal capsule polysaccharides are synthesized via a Wzx/Wzy-dependent pathway [4] , except for serotypes 3 and 37 which use a synthase-dependent pathway [4,5]. To date, 107 different pneumococcal serotypes have been defined based on their distinct capsular structures, serology, and antigenicity [6]. Based on the genetic sequences, Elberse *et al.* described genetic subtypes in a single serotype [7]. Importantly, genetic subtypes still form identical polysaccharide capsules within their respective serotypes and so are serologically identical to their respective serotypes.

The currently licensed vaccines against the pneumococcus target the polysaccharide capsule; the pneumococcal conjugate vaccines (PCVs) contain a subset of capsular polysaccharides conjugated to a carrier protein to improve efficacy [8], whilst the pneumococcal polysaccharide vaccine (PPV) contains pure pneumococcal polysaccharide [8]. However, each formulation only contains the polysaccharide for a subset of serotypes, and so does not offer protection against all serotypes of pneumococcus. The wide usage of these vaccines has caused shifts in serotype distribution globally due to the selective pressures applied by the vaccines on certain serotypes which can cause invasive pneumococcal disease (IPD); serotypes which are not covered by any PCV or PPV have become more common, whilst those covered by the vaccine have decreased in prevalence [9,10]. Therefore, the accurate classification of serotypes is critical for the reporting of virulent strains, informing future vaccine design, and evaluating vaccination implementation.

The serotype of a pneumococcal isolate has been traditionally determined using a laboratory method based on the reactions of the isolate to a defined set of antisera. This method is known as the Quellung reaction. This serotyping method requires lab experience and the reading of results could be subjective. With the advent of next generation sequencing, there has been an increasing use of bioinformatics tools to genetically infer the serotype of pneumococcal isolates. Several publicly available bioinformatics tools have been developed which utilise pneumococcal whole genome sequencing data to determine the serotype of the isolate. However, none of these tools are regularly updated. The most recently published tools PfaSTer [11] and PneumoKITy [12] have not been updated on their Github repository since 10^th^ March 2023 and 22^nd^ June 2022, respectively. Older tools such as PneumoCaT [13] and SeroCall [13,14] have not been updated since 2019. Since 2020, nine new pneumococcal serotypes have been discovered: 10D [15], 15D [16], 20C [17], 24C [18], 36A [19], 36B [19], 33E [20], 33G [21], 33H [22]. As these serotyping tools are not updated regularly, there is no bioinformatics tool that can identify these novel serotypes, and so isolates of that serotype cannot be accurately typed computationally. Therefore, potential changes in serotype distribution could be missed by these serotyping tools.

One of these bioinformatics tools is SeroBA [23]. SeroBA was published in 2018 and has not been updated since its publication. Hence, SeroBA is not able to identify the nine new serotypes described recently. SeroBA uses a k-mer (short nucleotide sequence) based method to identify the *cps* locus of an isolate from whole genome sequencing raw read data [23]. Then, SeroBA uses a combination of alleles, genes and mutations identified in the *cps* to predict serotypes with high concordance to phenotypic results and PneumoCaT [23]. The prediction of each serotype is based on a database containing reference *cps* sequences and genetic information that can be used to define and distinguish each serotype.

We created the SeroBAnk, which encompasses and visualises the *cps* region of all known pneumococcal serotypes for which there is a reference sequence of the *cps* locus and a capsular structure available. This resource is hosted on a publicly available dedicated webpage (https://www.pneumogen.net/gps/#/serobank). Building upon this resource, we developed SeroBA(v2.0), a tool that can accurately detect 102 of 107 known pneumococcal serotypes (except for serotypes 24B, 24C, 24F, 7D and 6H) and 18 genetic subtypes in serotypes 6A, 6B, 11, 19A, 19F, and 33F [7,20,24]. SeroBA(v2.0) can detect 9 more serotypes and 18 additional genetic subtypes than SeroBA(v1.0.7). We validated SeroBA(v2.0) on 26,306 genomes from the Global Pneumococcal Sequencing project, as well as simulated reads derived from reference *cps* sequences, and show that SeroBA(v2.0) can reliably detect the nine new serotypes as well as having high concordance (88.9%; 15,945/17,933) with phenotypic serotypes, and an even greater concordance with phenotypic serogroups (91.9%; 16,480/17,933) when available. Finally, we propose a community-contribution based approach to ensure that SeroBA(v2.0) stays maintained and updated as novel serotypes continue to be discovered. SeroBA(v2.0) can be accessed at https://github.com/GlobalPneumoSeq/seroba.

## Methods

### Developing the SeroBAnk

The SeroBAnk webpage summarises all 107 published serotypes, providing an atlas of the *cps* locus, serological profile, capsular structure [10] for each serotype, along with literature references and downloadable reference *cps* annotated sequences. The SeroBAnk webpage will be updated as novel serotypes are discovered. Additionally, the webpage also provides information on the formulations of 21 PCVs currently available on or under development and a timeline of pneumococcal vaccine history can be found at the bottom of the page.

#### Multi-Fasta Reference File

To create the multi-fasta reference file for the SeroBAnk, we identified all serotypes that had been discovered up until 1st August 2024 through literature review using keywords “pneumococci”, “serotype”, and “capsule” between 1st Jan 2020 to 1st Aug 2024, and the recent pneumococcal serotypes repository [6]. We also identified yet unpublished or soon-to-be published serotypes through ad-hoc consultation with pneumococcal experts. In cases where a reference sequence of the *cps* was available, we collated the reference sequences into a single file. However, in some cases only the raw read data were available instead of a reference sequence, and so we extracted the *cps* sequence manually. Shovill (v1.1.0) [25] was used to *de novo* assemble the reads and Bakta (v1.9.2) [26] to annotate the assembled genomes. By manually inspecting the annotation, it was possible to see where the *cps* region was in the genome. We then used samtools (v1.17) [27] to extract the *cps* region using the coordinates from the annotation. Finally, these *cps* sequences were also appended to the multi-fasta reference file in the SeroBAnk. Two serotypes (6H and 7D) [28,29], had no complete reference sequence or whole genome sequencing isolates available in the literature, and so it was not possible to extract the complete *cps* sequence.

We also included sequences for the known genetic subtypes within the serotypes 6A, 6B, 11F, 19A, 19F and 33F [5, 16, 24]. In total, there are 18 *cps* sequences which characterise the subtypes. These genetic subtypes are serologically identical to the respective serotypes by the Quellung reaction, and their capsular structures are currently unknown.

#### Genetic Information

We then gathered the genetic information which determined each serotype and subtype into a single .tsv file for the SeroBAnk, allowing for users to quickly identify the variants responsible for a particular serotype. In cases, there are well-defined genetic differences that can be used to determine a serotype, such as the presence or absence of specific genes, differing alleles for specific genes and mutations which code for amino acid sequence changes or truncations. However, in some cases there was no clear genetic basis to determine a serotype. For example, serogroup 24 shows inconsistent genetic patterns in serotypes 24B, 24C, and 24F - i.e. genetic heterogeneity within serotype 24B can lead to the same phenotype being expressed which can be identified by traditional methods such as Quellung [18]. Therefore, serogroup 24 isolates are either typed as 24A, which can be reliably identified through the absence of the *rbsF* gene [18], or 24B/24C/24F.

For a more detailed description of the genetic basis for each serotype and subtype, see **Supplementary data 1.**

#### Visualisation

For each serotype, the reference *cps* locus was visualised using Clinker (v0.0.29) [30]. The SeroBAnk data can also be visualised on Microreact at this link: https://microreact.org/project/seroba-v2.

### SeroBA(v2.0)

SeroBA(v2.0) was built upon SeroBA(v1.0.7) [23] by integrating the serotype-specific genes and mutations identified in SeroBAnk to enable detection of 102 of 107 known pneumococcal serotypes. SeroBA(v2.0) takes illumina next generation sequencing reads as input and produces an output file which includes two columns which show the serotype of the sample and the genetic subtype of the sample.

We used simulated reads and real-world data to validate SeroBA(v2.0). For the nine newly added serotypes, we used the “wgsim” utility of samtools (v1.17) [27] to simulate reads from the *cps* sequence of the reference genomes, and tested SeroBA(v2.0) on the simulated reads. To genetically predict the serotypes of real-world whole genome sequencing data, we ran SeroBA(v2.0) on 26,306 pneumococcal genomes in the Global Pneumococcal Sequencing Project (GPS) database. Most (17,933, 68.2%) had serologically-determined phenotypic serotypes by Quellung reaction or Latex agglutination. If whole genome sequenced (WGS) isolates with phenotypically determined serotypes were available in the literature, SeroBA(v2.0) was also tested using these isolates. We validated the subtypes in the same way.

Finally, we used the genetic serotyping predictions from SeroBA(v2.0) on the GPS Dataset to investigate the global distribution of the new serotypes. These genomes were sampled from 57 different countries, between 1989 and 2020. These published genomes passed quality control and were assembled as previously described for the global pneumococcal dataset [31,32]. The metadata for the isolates is available in **Supplementary Data 2**.

For the newly added serotypes or subtypes, we further validated the results by checking for the presence or absence of specific genes, alleles and mutations using an in-house script, independent of SeroBA(v2.0). This validation 100% supported the SeroBA(v2.0) calling. A more detailed description of the validation methodology can be accessed: https://github.com/Oliver-Lorenz-dev/seroba_validation?tab=readme-ov-file#methodology The test suite can be accessed at: https://github.com/Oliver-Lorenz-dev/seroba_validation. The final list of serotypes and genetic subtypes that can be typed by SeroBA(v2.0) can be found in **Supplementary Data 3**.

### SeroBA(v2.0) Concordance with Phenotypic Typing and PneumoCaT

To calculate the concordance of genetically predicted serotypes from SeroBA(v2.0) and phenotypically determined serotypes using Quellung/Latex agglutination, we investigated the genomes with a phenotypically determined serotype provided from the GPS database (Accessed 8th October 2024, available on Monocle https://data.monocle.sanger.ac.uk/) (n = 17,933). We then matched the genetically predicted serotyping results from SeroBA(v2.0) to the phenotypically determined serotyping results. We also used PneumoCaT (https://github.com/ukhsa-collaboration/PneumoCaT) to further investigate isolates for which the phenotypically determined serotypes and the genetically predicted serotypes from SeroBA(v2.0) were not concordant [13]. The concordance results can be found as **Supplementary Data 4,** with the PneumoCaT results alone in **Supplementary Data 5**.

## Results

### New serotypes detected by SeroBA(v2.0) 15D

Serogroup 15 previously contained four serotypes: 15F, 15A, 15B, and 15C. These can be genetically distinguished from each other; 15F diverges significantly from the other serotypes in this serogroup due to the presence of four additional genes: *glf*, *rmlB*, *rmlD* and *wcjE* (**Figure 1**) [33]. Serotypes 15B and 15C can be differentiated by a frameshift mutation in a tandem repeat in *wciZ* in 15C [34]. Serotypes 15B and 15C can be difficult to distinguish due to the high potential for a strain to interconvert between these two serotypes [35]. In addition, the quality of the local assembly must be high for SeroBA(v2.0) to differentiate these serotypes. If SeroBA(v2.0) is unable to differentiate these serotypes, it assigns a combined designation of 15B/15C. A novel serotype, 15D, was identified in 2021. Genetically, it is closest to 15A Serotype 15D can be distinguished from 15B and 15C due to *wzy15AF*, from 15A by its functional *wciZ*, and from 15F by its lack of *wcjE* [16] (**Figure 1**).

**Figure 1.**
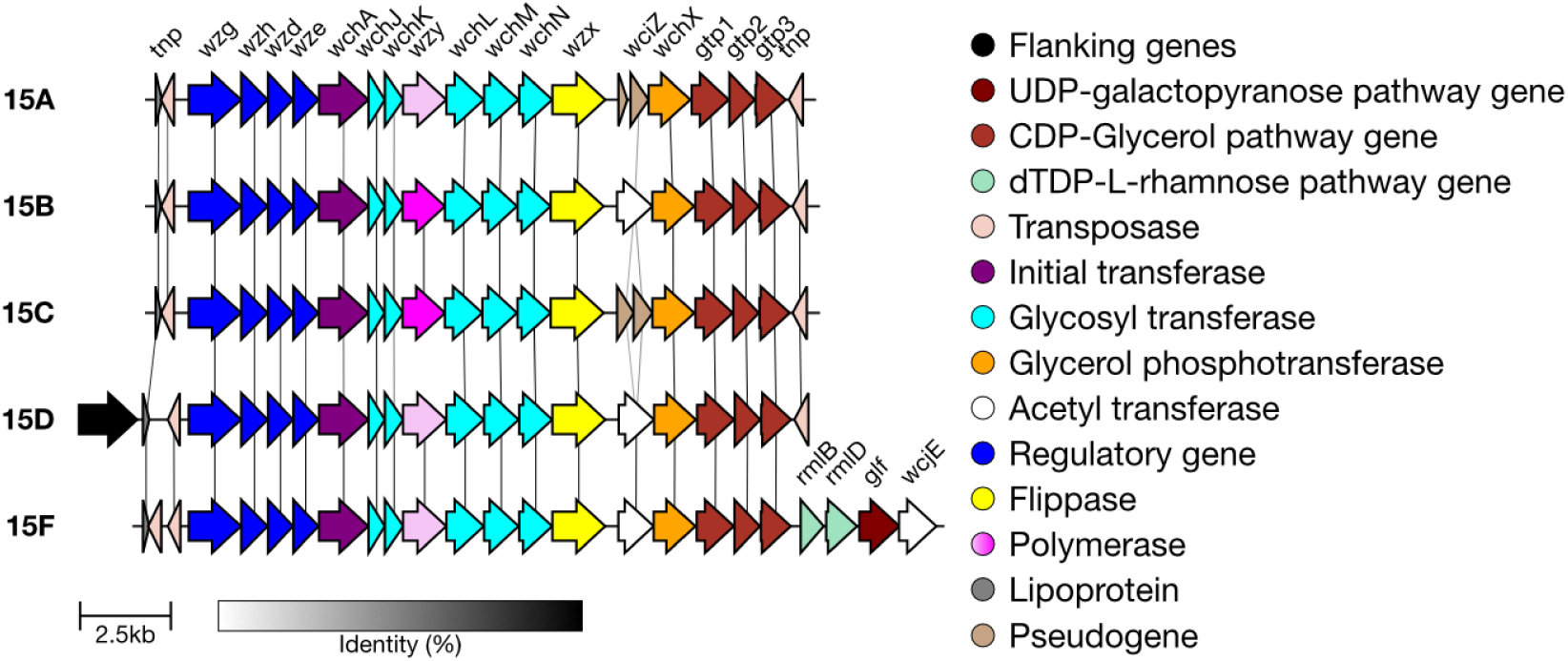
Comparison of the cps loci of serotypes in serogroup 15. Within serotype 15, there is the wzy15BC allele (dark pink) and the wzy15AF allele (pale pink). Serotype 15D can be distinguished from 15B and 15C due to wzy15AF, from 15A by its functional wciZ, and from 15F by its lack of wcjE [16].

Previously, SeroBA typed 15D isolates as either 15A or 15F. To validate the addition of serotype 15D to SeroBA(v2.0), we tested it by running SeroBA on a confirmed 15D WGS isolate sampled from the USA, SRR11130100 [16]. In addition, we used simulated reads derived from the 15D reference *cps* sequence (NCBI accession no. SRR11130100). Both were correctly typed as 15D. There were no further sequences in the GPS dataset that were typed as 15D. Serogroup 15 can be visualised on Microreact at https://microreact.org/project/seroba-serogroup15.

### Serogroup 20

Previously, SeroBA typed isolates belonging to serogroup 20 to only the serogroup level. However, the genetic basis of each serotype within serogroup 20 has since been identified. Serotype 20A is characterised by the loss of glycosyltransferase functionality of *whaF* [36], whilst serotype 20B has a functional *whaF* gene which encodes a glucose side chain, as compared with serotype 20A (**Figure 2**) [36]. The genetic basis of the more recently discovered 20C [17] is the loss of function of an O-acetyl transferase gene (*wciG*) in combination with the presence of the *whaF* gene (**Figure 2**). Interestingly, two isolates in the GPS database (GPS_US_PATH4317 and GPS_MW_C2307) have truncated *wciG* and *whaF* genes. These isolates could represent a novel serotype with capsular structure similar to 20A but absence of acetylation at the terminal galactofuranose.

**Figure 2.**
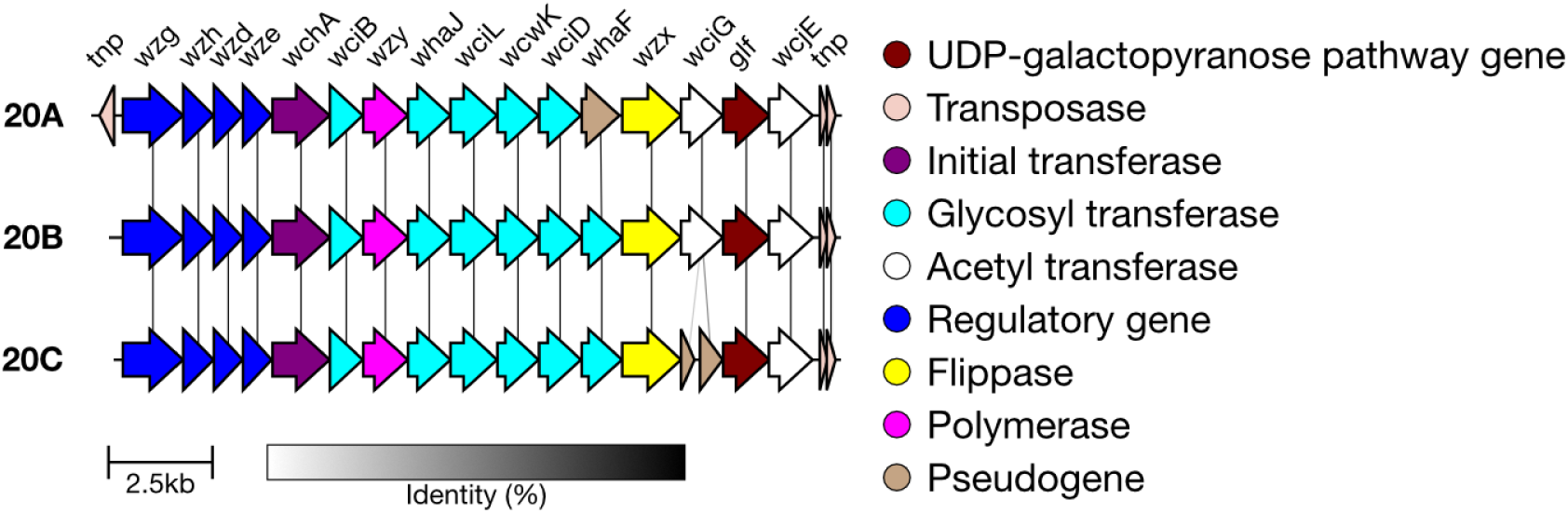
Comparison of the cps loci of serotypes in serogroup 20. Serotype 20A can be identified by the loss of function of whaF, and serotype 20C by the disruption of wciG.

Of the 167 serogroup 20 isolates in GPS, 123 are now typed as 20A, and 43 as 20B. One isolate had a “low coverage” error when run on SeroBA(v2.0) due to read quality issues, but manual inspection of the cps locus confirms it to be serotype 20C. Multiple read quality issues were observed for this sequencing run. This 20C isolate was from a carriage sample collected from an 8-month-old in 2014 in Nepal. It belongs to the lineage Global Pneumococcal Sequencing Cluster 43 (GPSC43). GPSCs are genomic definitions of pneumococcal lineages based on differences in the core and accessory genome [32]. We validated the addition of 20A, 20B and 20C using simulated reads from the *cps* reference sequences and real data. All simulated data were typed correctly. The validation scripts consistently identified the correct presence/absence of genes for each serotype in the WGS isolates. Test data output for <u>20A</u>, <u>20B</u> and <u>20C</u> are available for each serotype, and serogroup 20 can be visualised on Microreact at https://microreact.org/project/seroba-serogroup20.

### Serogroup 24

Previously, SeroBA also typed serogroup 24 isolates to the serogroup level only. It is possible to accurately type serotype 24A due to the lack of the *rbsF* gene compared to all other serotypes in this serogroup [18]. The loss of gene *rbsF* leads to a loss of ribofuranose biosynthesis in serotype 24A (**Figure 3**), resulting in absence of ribitol and ribofuranose in the polysaccharide structure [18]. The genetic basis of the other serotypes in serogroup 24 (24B, 24C and 24F) is not well defined, making it challenging to type isolates belonging to these serotypes with a high level of confidence [10].

**Figure 3.**
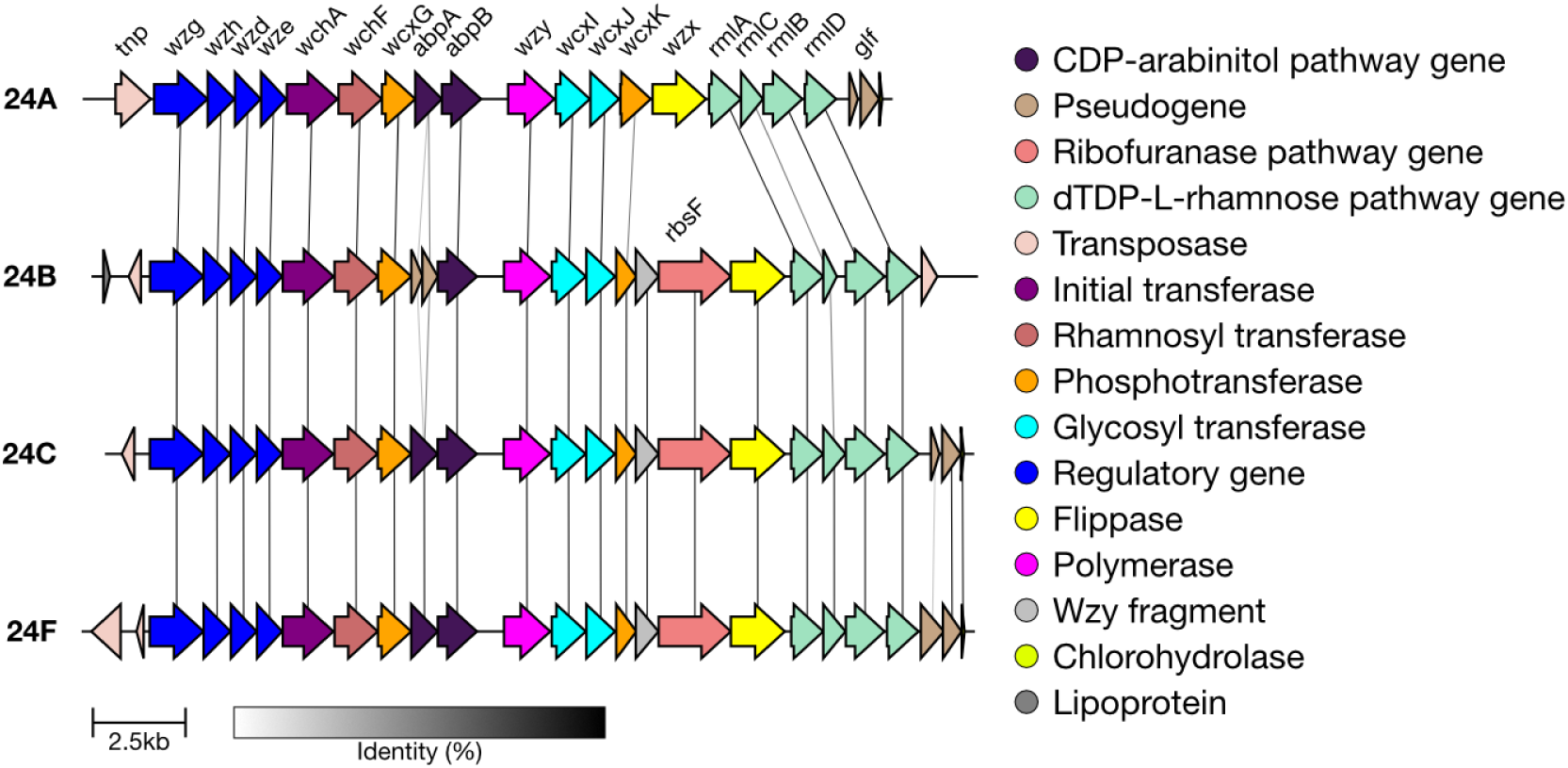
Comparison of the cps loci of serotypes in serogroup 24. Serotype 24A can be identified by the lack of gene rbsF. Although abpA is shown to have disruption in serotype 24B here, this disruptive mutation is not consistent among all 24B isolates observed so far (GPS database and 14), so this cannot be classified further by SeroBA(v2.0). Additionally, some 24B isolates do carry the glf pseudogene, which is not shown in this reference sequence [14].

Using SeroBA(v2.0), isolates belonging to serogroup 24 can now be typed as serotype 24A or 24B/24C/24F . Dissimilar to serotype 24F, serotype 24B lacks arabinitol in the repeat unit structure, probably due to the disruptive function of *abpA*. However, disruptive mutation in *abpA* was not consistently observed among the phenotypic serotype 24B isolates (n=9), which were typed as serogroup 24 by SeroBA(v1.0.7) in the GPS database. It is also of note that a German serotype 24B isolate MNY583 had an intact AbpA with D222Y nonsynonymous substitution reported in previous study [18]. It is possible that substitution(s) in *abpA* may potentially affect the enzymatic function, rather than by disruptive mutation, making it challenging to differentiate serotypes 24B and 24F. In addition, serotype 24C is a mixture of serotype 24B and 24F with ratios varying between strains [18]. Considering the heterogeneous genetic basis observed within each serotype, SeroBA(v2.0) grouped 24B/24C/24F in the output until further experimental validation.

We tested the serotyping of 24A using simulated reads derived from reference *cps* sequence (NCBI accession no. CR931686) and isolates which were phenotypically typed as 24A (n = 4) in the GPS dataset. The test was 100% concordant. In the GPS dataset, there are 670 sequences belonging to serogroup 24. Of these sequences, 11 were typed as 24A whilst the rest were typed as 24B/24C/24F. We validated this result using a test script which confirmed none of these isolates contained the *rbsF* gene. Test data output are available for <u>24A</u>. The 11 24A sequences were collected between 2001-2017 in India (n = 3), Malawi (n = 3), Nepal (n = 2), Nigeria (n = 1), South Africa (n = 1) and Thailand (n = 1). Most isolates belong to GPSC91 (81.8%; n = 9), however the Thai sample belonged to GPSC106 and the South African sample belonged to GPSC966. The genomes can be visualised on Microreact at this link: https://microreact.org/project/seroba-serogroup24.

## 33E

Serotype 33E was previously typed as 33F by SeroBA. The genetic basis of serotype 33E is characterised by the inactivation of *wciE* that encodes glycosyltransferase which transfers glucose due to a nonsense mutation [20] (**Figure 4**). The inactivation of *wciE* causes a lack of glycosyltransferase activity in 33E, which gives the capsular polysaccharide a unique repeat unit backbone structure with no branching that does not contain ribitol [10,20].

**Figure 4.**
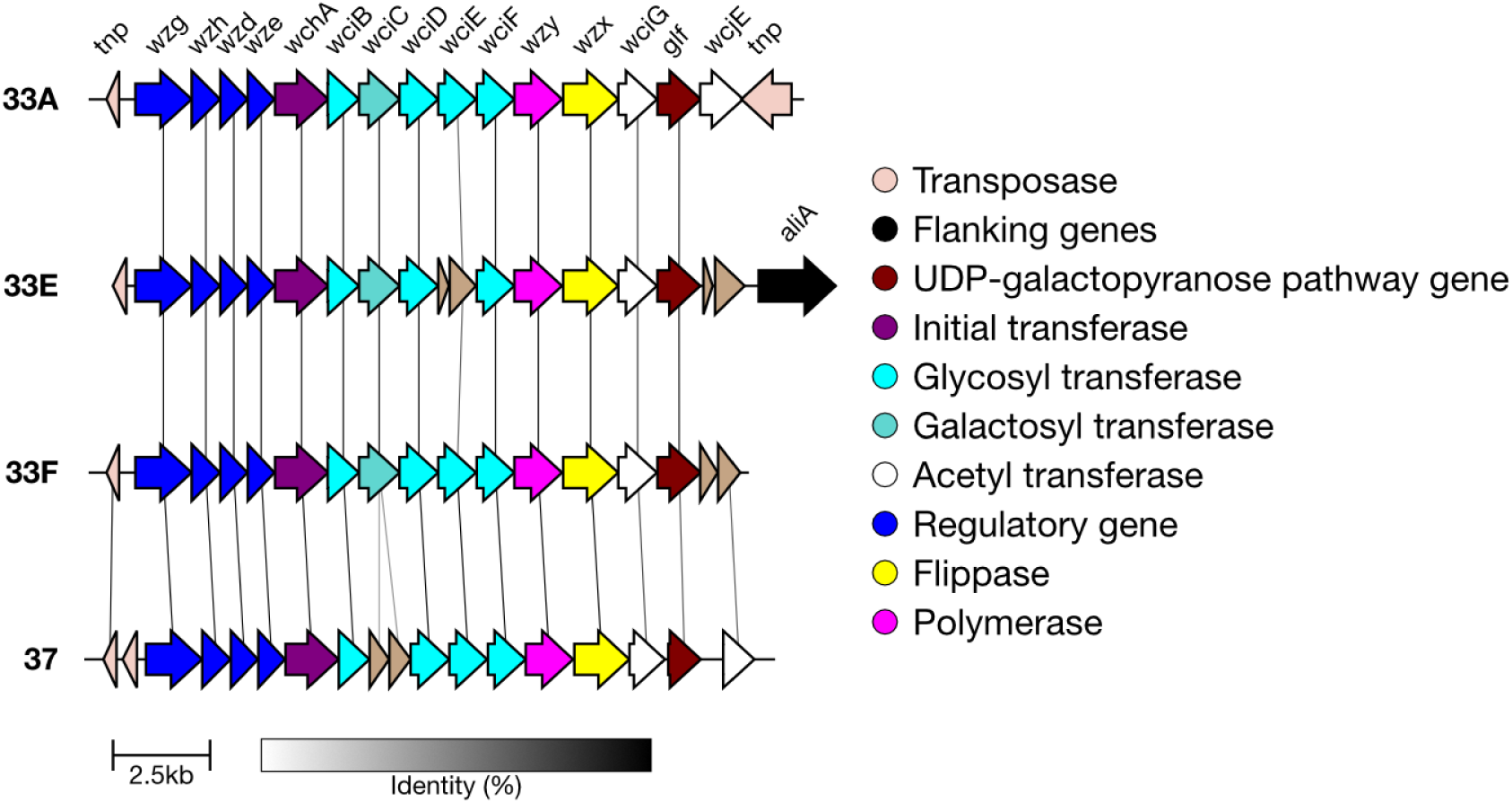
Comparison of the cps loci of serotype 33E and closely related serotypes. Serotype 33E can be identified due to the inactivation of wciE. Serotype 37 can be differentiated due to a glucosyl transferase gene called tts, found outside of the cps region. tts is responsible for the polysaccharide structure of serotype 37, where the repeat unit consists of two glucopyranose molecules connected by a β1-2 linkage [5,10].

We validated the serotyping of 33E using simulated reads derived from 33E reference *cps* sequences (NCBI accession no. SAMEA2203953) and two isolates from the GPS dataset which were phenotypically identified as 33E [20]. The test was 100% concordant, with SeroBA(v2.0) correctly identifying them as serotype 33E. Test data output is available for <u>33E</u>. These isolates were both from paediatric patients with IPD, a <1 year old in the USA, and a 10-month-old in New Zealand. Both are also members of the lineage GPSC3. No other isolates were identified as 33E in the GPS dataset. Serogroup 33 can be visualised on Microreact at https://microreact.org/project/seroba-serogroup33.

### 33G and 33H

SeroBA(v2.0) can type both serotypes 33G [21] and 33H [37], yet were both typed as 10X by SeroBA(v1.0.7). SeroBA uses k-mer (short nucleotides) identity to distinguish 33G and 33H from other serogroup 33 members. The genetic basis for distinguishing 33H from 33G is a premature stop codon that inactivates the acetyltransferase *wciG* (**Figure 5**). The Quellung reactions of 33G and 33H show as 10B+33B and 10B+33F, respectively [21,37], and the 11 serotype 33G isolates were typed as 10F (n=8), followed by 33F (n=1); two isolates were not typed by Quellung.

**Figure 5.**
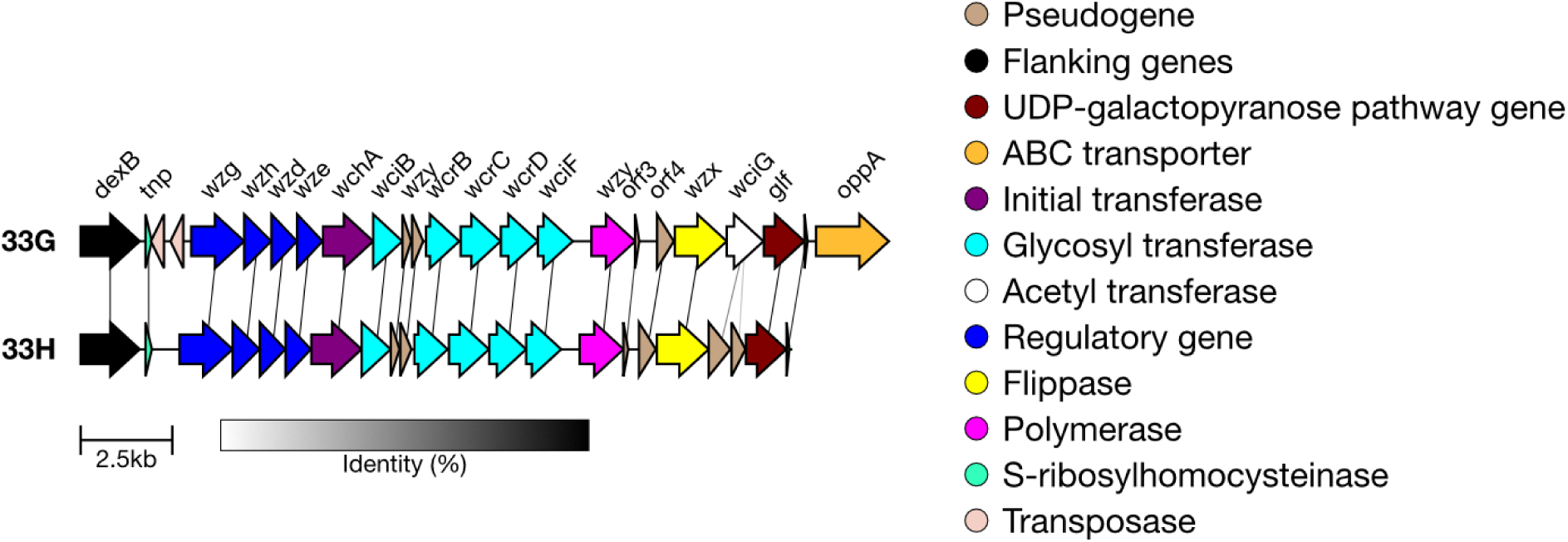
Comparison of the cps loci of serotype 33G and 33H. Serotype 33H can be identified due to the inactivation of wciG due to a premature stop codon.

We tested the genetically predicted serotyping of SeroBA(v2.0) on serotype 33H using one isolate from the GPS dataset which had an inactivated *wciG*. We validated that *wciG* was inactivated in this sample. Test output are available for <u>33H</u>. This was the only 33H isolate in the GPS dataset, which was isolated in South Africa in 2007 from a case of IPD in a child <5 years of age, which was phenotypically typed as 10B by Quellung. Serogroup 33, including 33G and 33H can be visualised on Microreact at https://microreact.org/project/seroba-serogroup33.

### Serogroup 36

Previously, SeroBA typed serogroup 36 isolates at the serogroup level only. Serotype 36B was recently discovered and differs from 36A due to a divergent *wcjA* allele (S144G, F269M, G270A, A273V) which has differing glycosyltransferase activity (**Figure 6**) [19]. Because of this, there is a distinguishing glycosidic bond in the polysaccharide which incorporates Galp in 36A versus Glcp in 36B [19].

**Figure 6.**
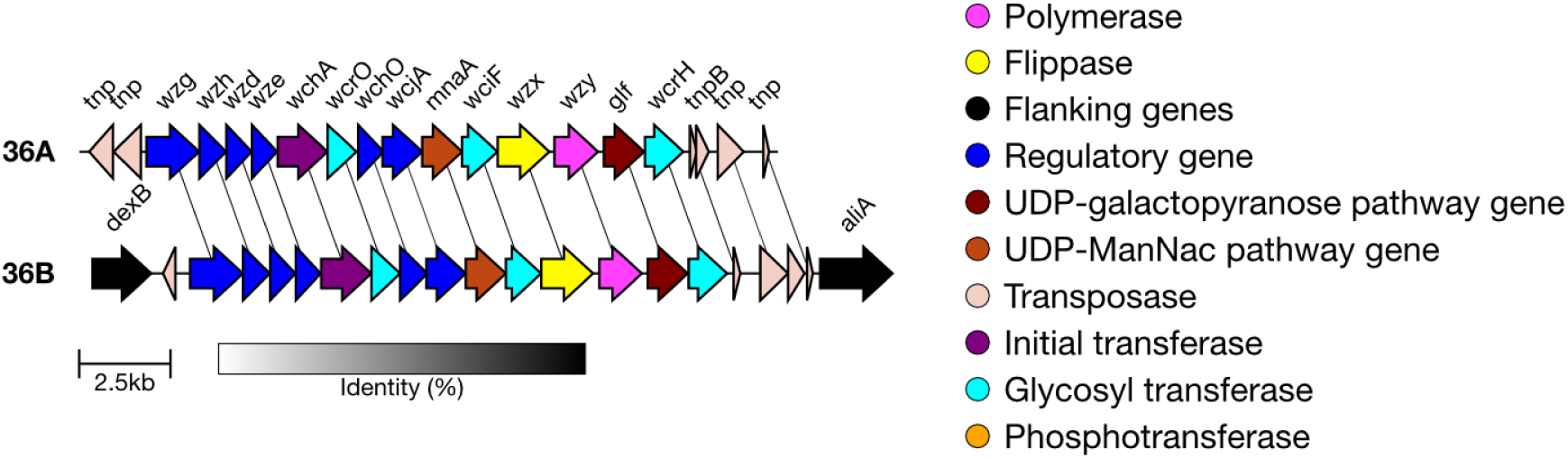
Comparison of the cps loci of serotype 36A and 36B. The two can be differentiated by divergent wcjA alleles.

We validated the serotyping of 36B using simulated reads derived from the *cps* reference sequences and the nine serogroup 36 sequences in the GPS dataset. Test data output is available for <u>36A</u> and <u>36B</u>. Three of these isolates were typed as 36B, two of which were confirmed as serotype 36B by serology using monoclonal antibodies and capsular structure using NMR by Ganaie et al [19]. The other sample was GPS_IN_CS293, which was collected from a patient with IPD in India in 2005. The sample belongs to pneumococcal lineage GPSC241, ST15709. Serogroup 36 can be visualised on Microreact at https://microreact.org/project/seroba-serogroup36.

### New Genetic Subtypes

Genetic subtypes of serogroups 6 and 19 were obtained from [7], 11F from [24] and 33F from [20].

### 11F-like (also known as 11X)

Genetic subtype 11F-like isolates (also known as 11X [38]) have previously been identified in Fiji and can be typed using microarray-based methods [24]. The capsular structure of the 11F-like *cps* is identical to that of 11A; isolates are typed as 11A by the Quellung reaction, despite being genetically more similar to serotype 11F. Previously, SeroBA would type 11F-like isolates as 11A.

We found that the key genetic differences which characterise the genetic variant 11F-like are two divergent alleles of *wcrL* (S17P, A112N, and D165N) and *wcwC* (Multiple nonsynonymous substitutions, 90.9% amino acid identity) (**Figure 7**) (**Supplementary Data 1**). The A112N mutation in *wcrL* means that the protein transfers αGlc, as opposed to αGlcNAc, and so the 11F-like capsule would lack this modification [24]. Additionally, unlike in 11F, the *gct* gene is intact in 11F-like. *gct* encodes for a CDP-glycerol synthetase, which catalyses the biosynthesis of Glycerol-1-phosphate (Gro1p), and so we would expect Gro1p to be present in the 11F-like capsule [24].

**Figure 7.**
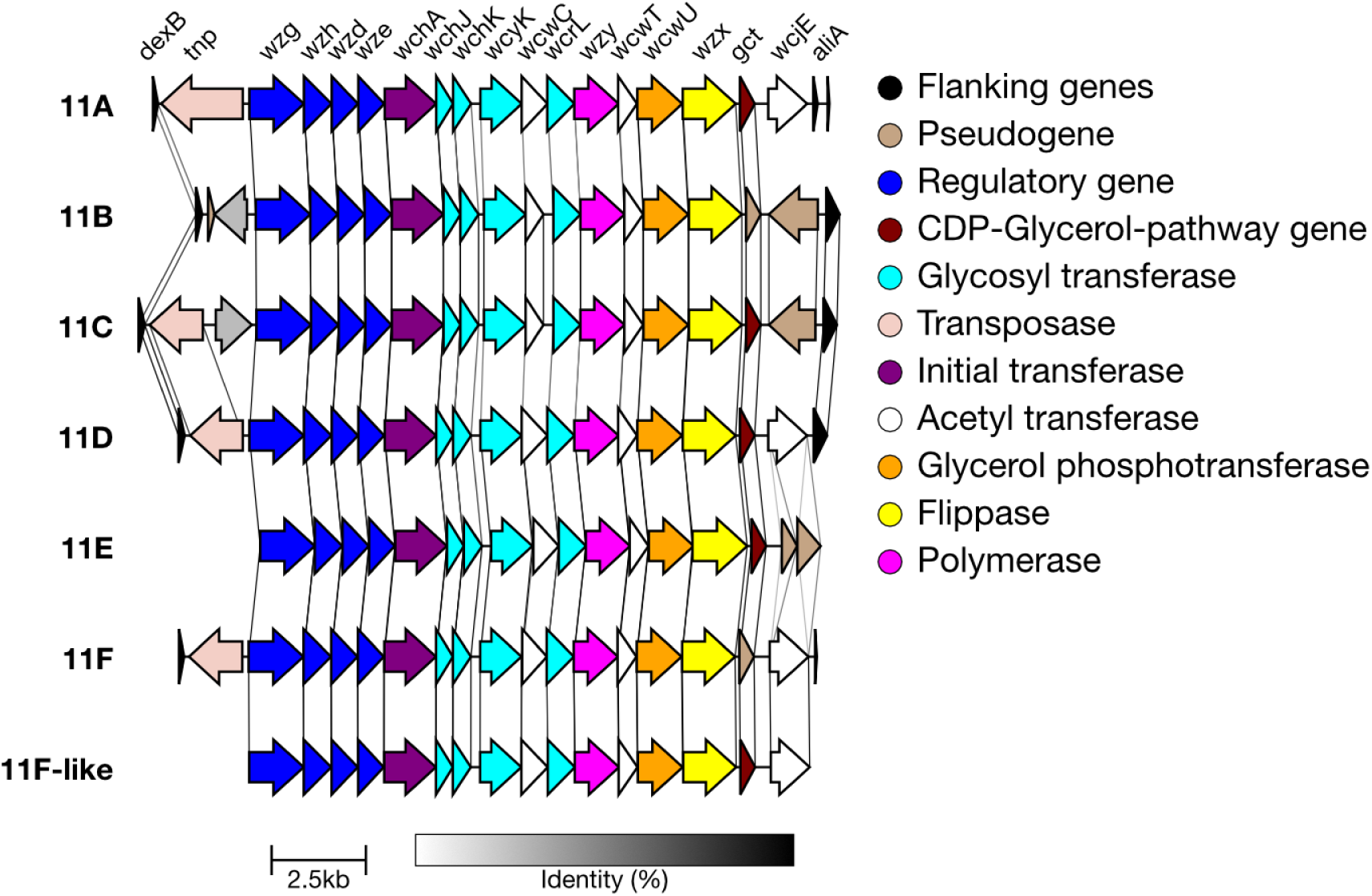
Comparison of the cps loci of serotypes in serogroup 11. The new genetic subtype 11F-like can be differentiated from serotype 11F and 11B due to activity of the gct gene, whilst it can be distinguished from 11A, 11C, 11D, and 11E by differing wcwC and wcrL alleles.

Using SeroBA(v2.0), 52 isolates from the GPS dataset were typed as the genetic variant 11F-like, where they had previously been typed as 11A. We validated that these sequences contained the correct genes, and test data output are available for <u>11F-like</u>. The genomes were collected between 2000 and 2017 and were mainly located in Asia. 76.9% of 11F-like isolates were from Cambodia (GPSC6, n = 30; GPSC10, n = 1; GPSC73, n = 8; GPSC626, n = 1), 19.2% from China (GPSC73, n = 9; GPSC6, n = 1), and a single sample each from Ghana (GPSC360) and South Africa (GPSC360). These data can be viewed at https://microreact.org/project/seroba-serogroup11.

### 6A genetic subtypes

In addition to the reference type, there are six genetic subtypes defined within serotype 6A: 6A-I, 6A-II, 6A-III, 6A-IV, 6A-V and 6A-VI [7], each varying to a different degree from the reference serotype 6A. Subtype 6A-II has the most amino acid substitutions compared to the 6A reference sequence, whilst 6A-IV has one [39]. 6A-IV can be identified by a non-synonymous mutation (A163V) in the *wze* gene. The other five subtypes each have at least two divergent alleles which include non-synonymous mutations compared to the 6A reference sequence. The divergent alleles can be seen in genes including *wzg*, *wzy*, *rmlA*, *rmlB* and *rmlC* (**Figure 8**).

**Figure 8.**
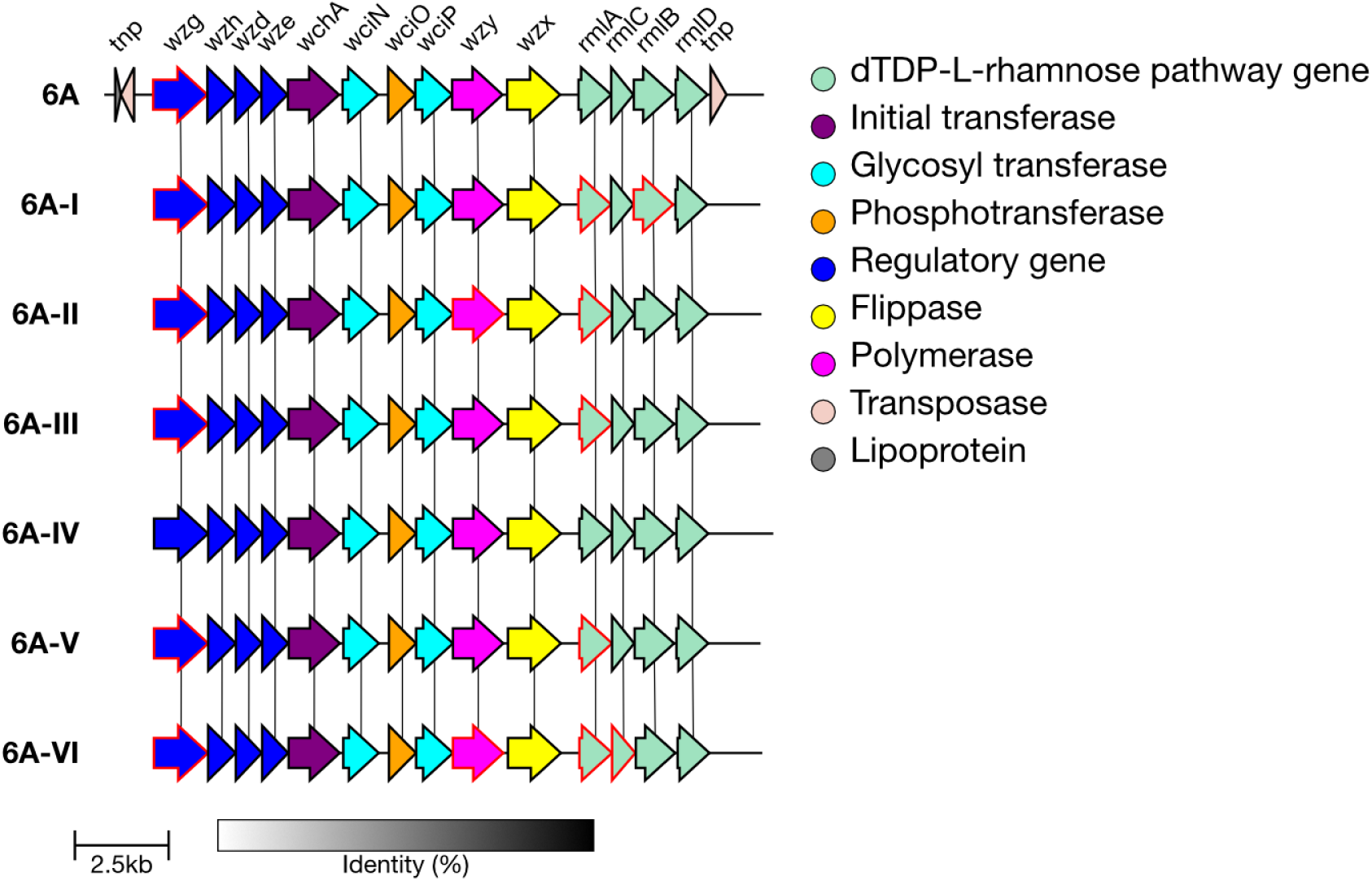
Comparison of the cps loci of serotype 6A genetic subtypes. Genes which have 10 or more base pairs different from the 6A reference sequence are outlined in red. 6A-IV can be distinguished by a single non-synonymous mutation in wze.

**Figure 9.**
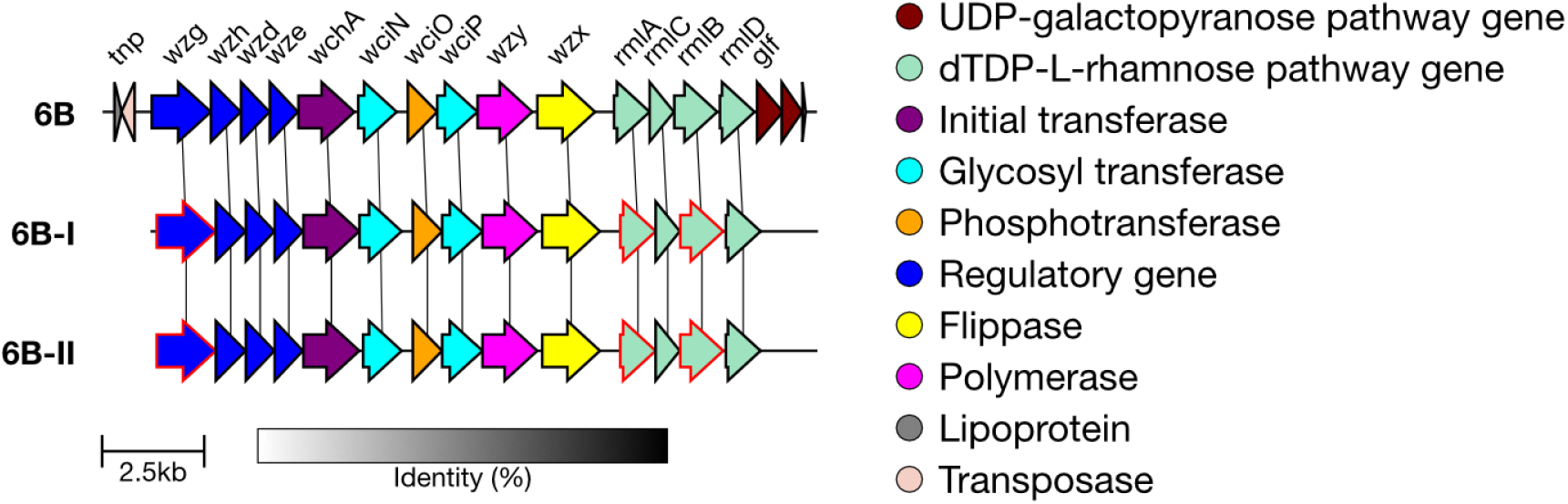
Comparison of the cps loci of serotype 6B genetic subtypes. Genes which have 10 or more base pairs different from the 6B reference sequence are outlined in red.

**Figure 10.**
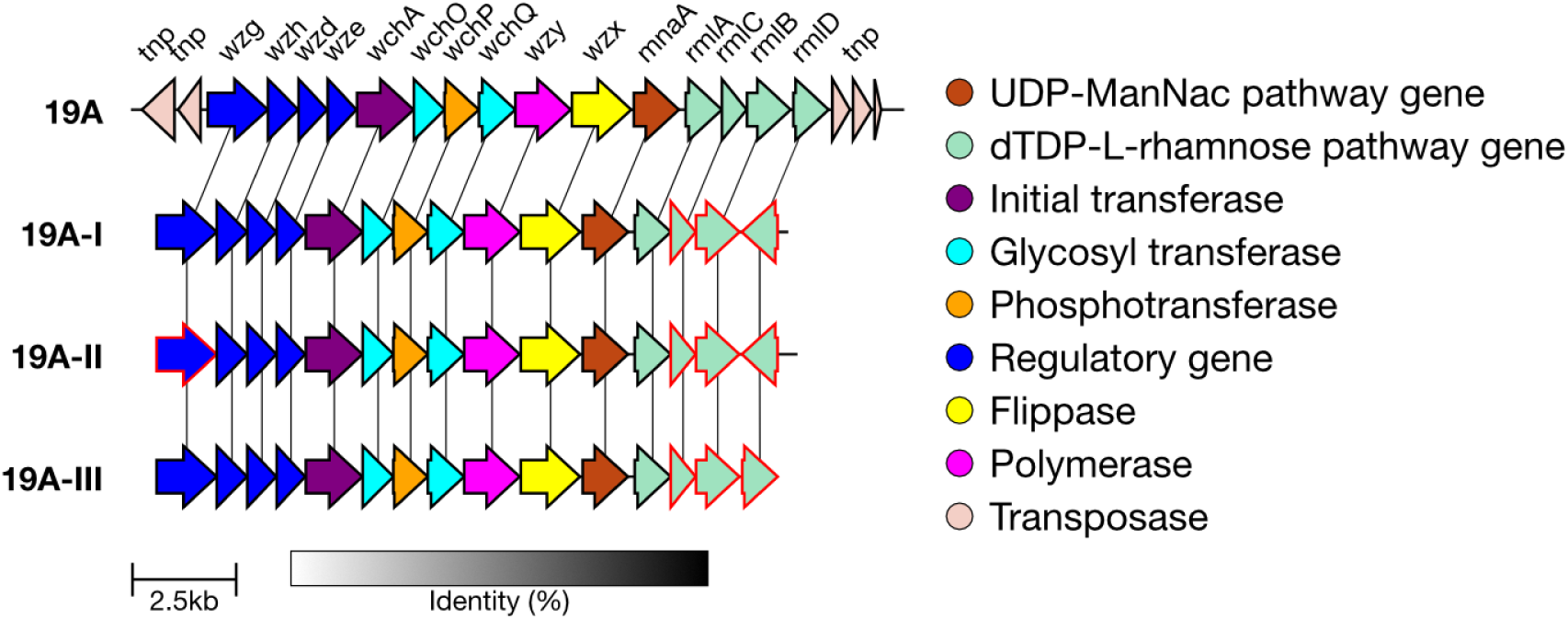
Comparison of the cps loci of serotype 19A genetic subtypes. Genes which have 10 or more base pairs different from the 19A reference sequence are outlined in red.

**Figure 11.**
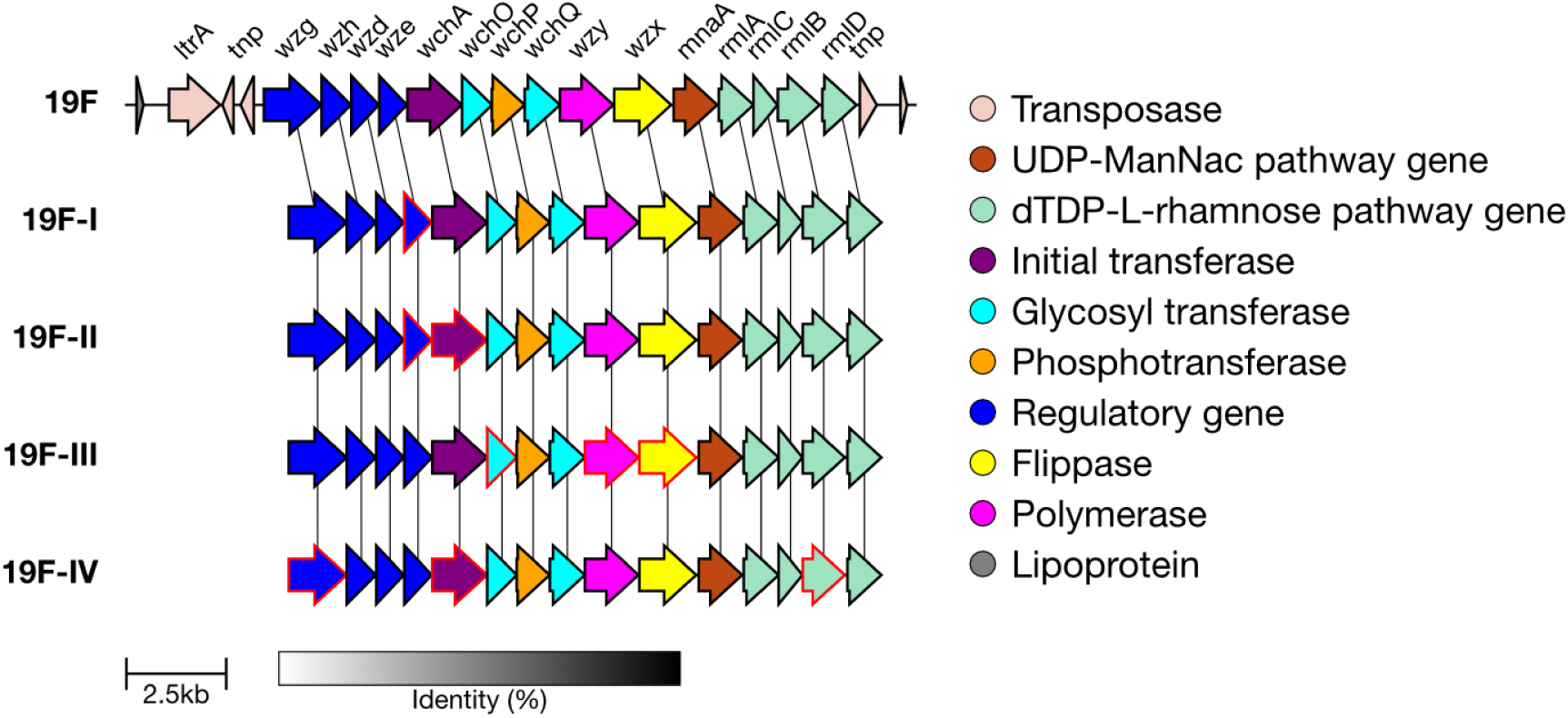
Comparison of the cps loci of serotype 19F genetic subtypes. Genes which have 10 or more base pairs different from the 19F reference sequence are outlined in red.

In the GPS dataset, 1,603 isolates were typed as 6A by SeroBA(v2.0). Of these, 960 were typed as the 6A reference sequence, 400 were typed as 6A-II, 212 as 6A-I, 18 as 6A-III, 9 as 6A-V, 3 as 6A-VI and 1 as 6A-IV. This highlights the high level of genetic diversity within this serotype. A Microreact showing these isolates is available at https://microreact.org/project/seroba-serogroup6.

We validated that the correct alleles or mutations were present in a randomly selected subset of sequences (maximum 100) for each subtype. Test data outputs are available for <u>6A-I</u>, <u>6A-II</u>, <u>6A-III</u>, <u>6A-IV</u>, <u>6A-V</u> and <u>6A-VI</u>.

### 6B genetic subtypes

In addition to the reference type, there are two genetic subtypes defined for serotype 6B: 6B-I and 6B-II [7]. Both of these subtypes are characterised by three divergent alleles within *wzg*, *rmlA* and *rmlB*, which have non-synonymous mutations compared to the 6B reference sequence (**Supplementary Table 1**). The alleles seen within 6B-I and 6B-II are different, allowing for the two subtypes to be distinguished.

In the GPS dataset, 217 isolates were typed as 6B by SeroBA. Of those, 111 were typed as the 6B reference sequence, 102 were typed as 6B-I and 4 as 6B-II. These can be seen on Microreact at https://microreact.org/project/seroba-serogroup6.

We validated that the correct alleles or mutations were present in a randomly selected subset of sequences (maximum 100) for each subtype. Test data outputs are available for <u>6B-I</u> and <u>6B-II</u>.

### 19A genetic subtypes

In addition to the reference type, there are three genetic subtypes defined for serotype 19A: 19A-I, 19A-II and 19A-III [7]. These subtypes are characterised by divergent alleles in the rhamnose related genes – *rmlB*, *rmlC* and *rmlD*. All three subtypes contain an identical *rmlC* allele which diverges from the reference *rmlC* allele by 51 bases in the final 231 bases of the gene.

19A-I and 19A-II both have divergent *rmlB* alleles which differ from the reference by over 160 bases, and a highly divergent *rmlD* allele which is oriented in the opposite direction to *rmlD* in the reference sequence. However, 19A-I and 19A-II can be distinguished due to a single non-synonymous mutation (A143S) in the *wzg* gene. If SeroBA(v2.0) is unable to detect this mutation, but the divergent *rmlB*, *rmlC*, and *rmlD* alleles are all detected, it will type the sample as 19A-I/19A-II, as it cannot be confident of either result. 19A-III also has divergent *rmlB* and *rmlD* alleles, but these are closer to the reference sequence than those of 19A-I/19A-II, and so it can be identified.

We validated that the correct alleles or mutations were present in a randomly selected subset of sequences (maximum 100) for each subtype. Test data outputs are available for <u>19A-I</u>, <u>19A-II</u> and <u>19A-III</u>.

In the GPS dataset, 2,136 isolates were typed as 19A by SeroBA(v2.0). Of those, 968 were typed as the 19A reference sequence, 877 were typed as 19A-I/19A-II, 75 as 19A-I, 215 as 19A-II and one as 19A-III. These data can be seen on Microreact at https://microreact.org/project/seroba-serogroup19.

### 19F genetic subtypes

In addition to the reference type, there are four genetic subtypes defined for serotype 19F: 19F-I, 19F-II, 19F-III and 19F-IV [7]. 19F-I is characterised by SNPs in *wze* (E45R, I159), and 19F-II also has SNPs in *wze* alongside a divergent *wchA* allele. 19F-III is similar to the 19F reference, though it has a small number of non-synonymous mutations in *wchO* (S22I), *wzx* (I370M, I377L) and *wzy* (I71L) that allow it to be identified. 19F-IV diverges the most from the 19F reference - it has divergent *wchA*, *wzg* and *rmlB* alleles with amino acid similarity at 94.5%, 96.2% and 98.2%, respectively.

We validated that the correct alleles or mutations were present in a randomly selected subset of sequences (maximum 100) for each subtype. Test data output are available for <u>19F-II</u>, <u>19F-III</u> and <u>19F-IV</u> (19F-I was not typed using any GPS data but was typed correctly with simulated reads derived from the CPS reference sequence).

1,827 isolates in the GPS dataset were typed as 19F by SeroBA. Of those, 1,443 were typed as the 19F reference sequence, 223 as 19F-II, 160 as 19F-III, 1 as 19F-IV and 0 as 19F-I. These can all be seen on Microreact at https://microreact.org/project/seroba-serogroup19.

### 33F genetic subtypes

In addition to the 33F reference type, there are two genetic subtypes defined for serotype 33F: 33F-1a and 33F-1b [20]. 33F-1a differs from the 33F reference sequence by having an intact *wcyO* gene (an acetyl-transferase) instead of the *wcjE* pseudogene [20]. 33F-1b also has the *wcyO* gene instead of the *wcjE* pseudogene, however it is not intact [20].

We validated that the correct genetic combinations were being identified using typing results from [20]. Ganaie *et al.* evaluated 18 33F-1a and 10 33F-1b isolates from the GPS dataset. SeroBA(v2.0) was able to correctly identify all of these isolates, aside from a 33F-1a isolate (GPS_IL_26764) which was typed as 33A/33E/33F (undetermined) https://microreact.org/project/seroba-serogroup33.

Overall, 141 isolates in the GPS dataset were typed as 33F by SeroBA(v2.0). Of these, 100 were typed as the 33F reference sequence, 26 were typed as 33F-1a and 16 were typed as 33F-1b.

### Additional Non-encapsulated References

Two non-encapsulated reference sequences were in the SeroBA(v1.0.7) reference database. These were named “Swiss_NT” and “alternative_aliB_NT”. These reference sequences were renamed to provide a more detailed description of what each sequence is: “Swiss_NT” is now “NCC2_aliC_aliD_non_encapsulated” [40,41] and “alternative_aliB_NT” is now “NCC2_S_mitis_aliC_aliD_non_encapsulated” [40,41], to better reflect the gene content that characterises the two variants. Two additional non-encapsulated reference sequences have been added: “NCC3_aliD_non_encapsulated” which is similar to NCC2 except for the absence of *aliC* [40], and “NCC1_pspK_non_encapsulated”, which has the *pspK* gene [40]. *pspK* is known to compensate for the loss of virulence in non-encapsulated pneumococci [42].

269 isolates in the GPS dataset were typed as “Swiss_NT” by SeroBA(v1.0.7). These are now typed as “NCC2_aliC_aliD_non_encapsulated”. 3 isolates in the GPS dataset were typed as “alternative_aliB_NT” by SeroBA(v1.0.7), and these are now typed as “NCC2_S_mitis_aliC_aliD_non_encapsulated”. 95 isolates in the GPS dataset were typed as “NCC1_pspK_non_encapsulated” by SeroBA(v2.0). 94 of these isolates were determined to be “untypable” by SeroBA(v1.0.7). One “NCC1_pspK_non_encapsulated” isolate was typed as 23F by SeroBA(v1.0.7), we manually investigated the *cps* region of this isolate and found that whilst some of the transposes found within the 23F *cps* region were present, there was not an intact 23F *cps* region. The addition of NCC1_pspK_non_encapsulated is a better match, leading to the reassignment in SeroBA(v2.0). No isolates in the GPS dataset were typed as “NCC3_aliD_non_encapsulated”. The non encapsulated isolates can be visualized on Microreact at https://microreact.org/project/seroba-non-encapsulated. These non encapsulated isolates are described in Figure 13.

### Validation Against Phenotypic Results

New serotypes and genetic subtypes added to SeroBA(v2.0) were validated using reference isolates, data from the GPS dataset and simulated reads from the *cps* reference sequences. In addition, we further validated whether the SeroBA(v2.0) update impacted its accuracy for serotypes already within the software. For this, we compared the genetic serotype predictions of SeroBA(v2.0) to the phenotypically determined serotyping results from the GPS dataset. Of the 26,306 isolates within the dataset, 17,933 (68.2%) had a phenotypic serotype determined by serological methods (Quellung reaction or latex agglutination) (**Figure 12**). We found that 92.3% (n = 16,544/17,933) matched at either the serogroup or serotype level, with 89.3% (n = 16,009/17,933) matched at the serotype level.

**Figure 12.**
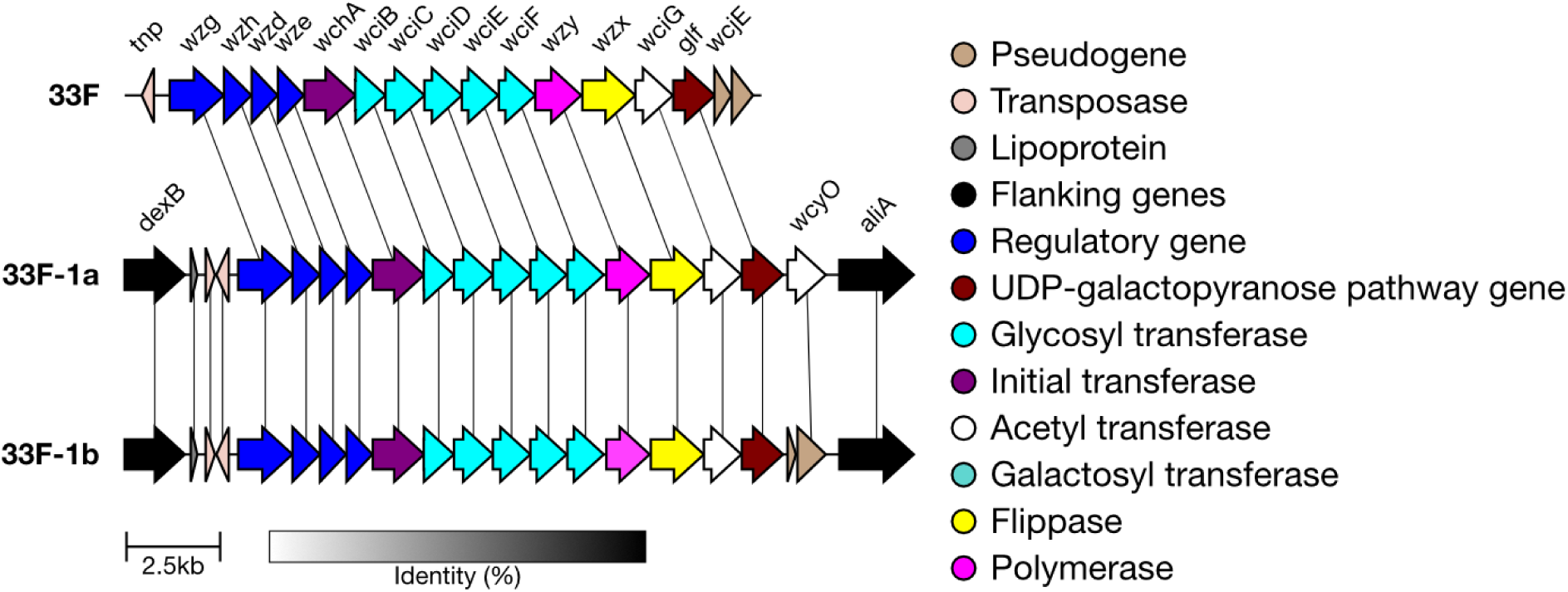
Comparison of the cps loci of serotype 33F genetic subtypes. 33F-1a contains a functional wcyO gene, whilst 33F-1b contains a non-functional wcyO gene.

**Figure 13.**
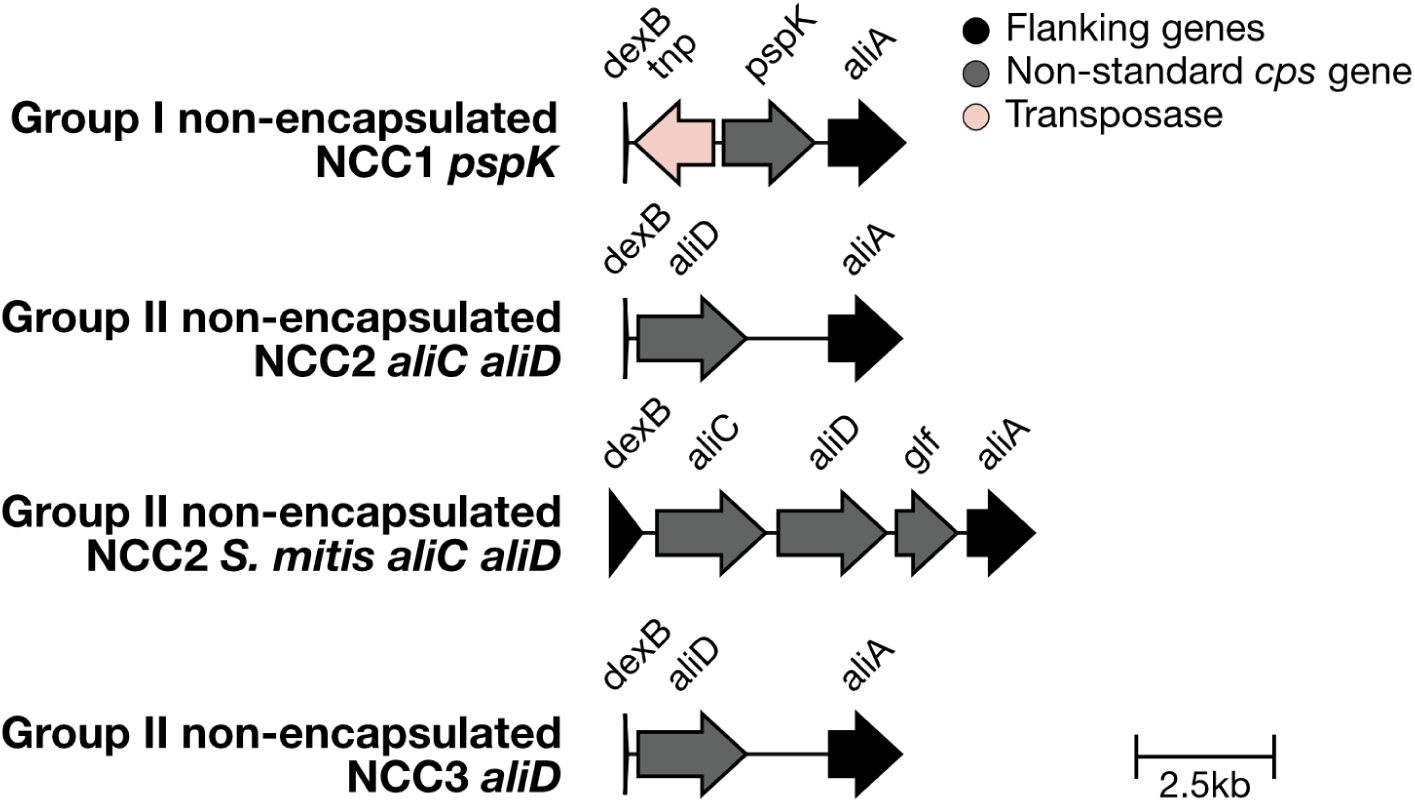
**Additional non encapsulated reference sequences in the SeroBA(v2.0) database.**

**Figure 13.**
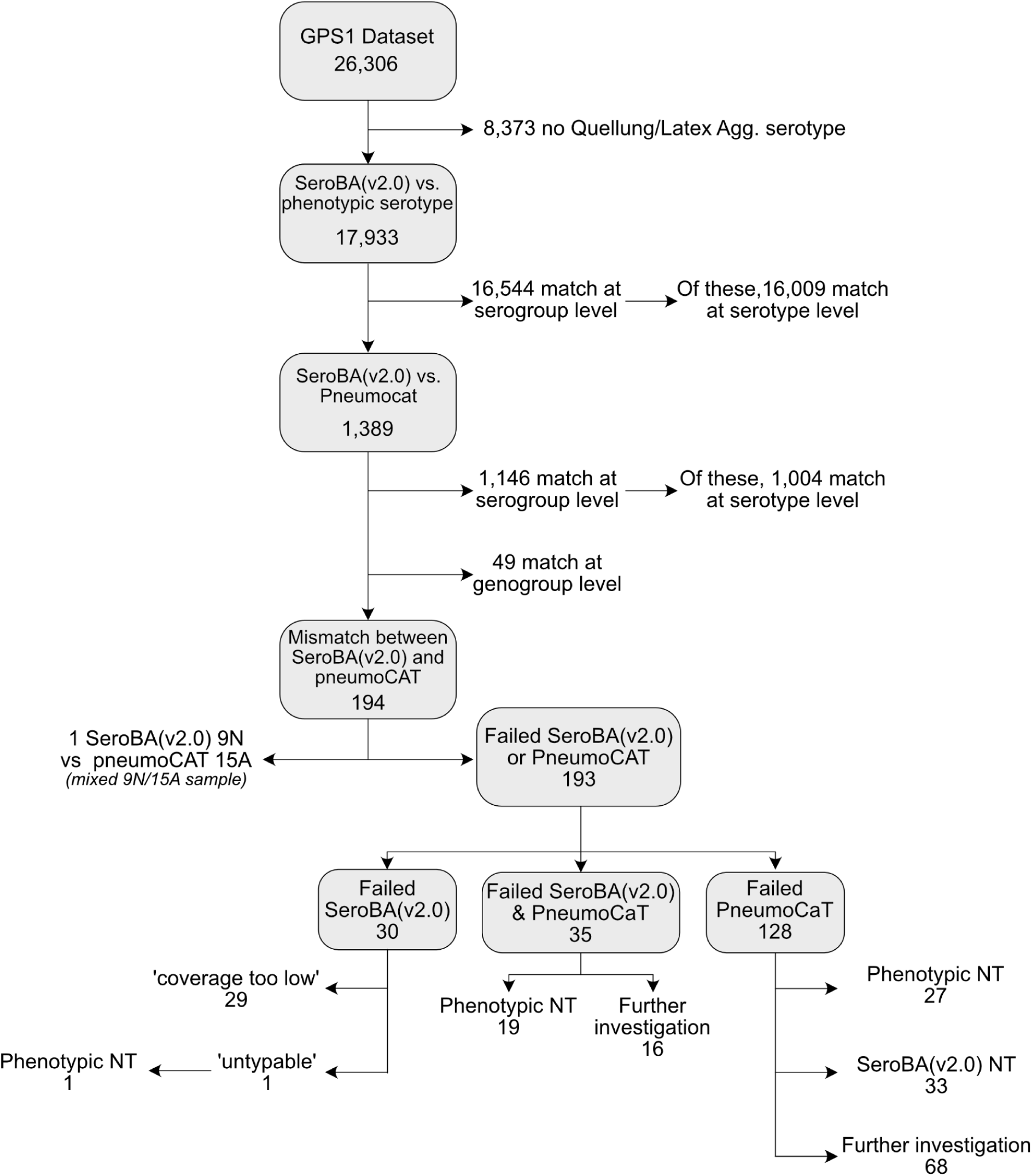
Validation of SeroBA(v2.0). The vast majority of the GPS database was concordant with SeroBA(v2.0) or PneumoCaT results, either at the serotype or serogroup level. PneumoCaT Genogroups are defined as groups of serotypes with >90% sequence identity within the cps locus [13].

We further investigated the isolates that did not match at the serogroup level [33]. We serotyped the remaining 1,389 isolates using the bioinformatics tool PneumoCaT [10] and compared the concordance with SeroBA(v2.0). 82.5% of isolates with SeroBA(v2.0)/phenotype mismatches were concordant between SeroBA(v2.0) and PneumoCaT at the serogroup level (n = 1,146/1,389) (**Figure 12**). Of the remaining 243 sequences, 49 matched between SeroBA(v2.0) and PneumoCaT at the genogroup level (e.g. serotypes 7B and 40, serotypes 12A and 46), defined as groups of serotypes with >90% identity in their *cps* locus [13]. One isolate (GPS_ZA_2871) was typed as 9N by SeroBA(v2.0) and 15A PneumoCaT and phenotype, however upon further investigation it was found to be a mixed sample containing both of these serotypes (**Figure 12**). The remaining 193 isolates failed their SeroBA(v2.0) run and/or PneumoCaT run (**Figure 13**).

128 isolates had a successful SeroBA(v2.0) run, but failed their PneumoCaT run (**Figure 13**). One cause of this can be that the isolate is non-encapsulated; in line with this, we found that 46.9% of isolates that failed PneumoCaT (n = 60/128) were either phenotypically non-typable or SeroBA(2.0) typed them as non-encapsulated strains.

35 isolates failed both their SeroBA(v2.0) and PneumoCaT runs (**Figure 13**). 54.3% (n = 19/35) of these were phenotypically non-typable, thus these isolates are likely non-encapsulated.

Finally, 30 isolates failed their SeroBA(v2.0) run, of which 96.7% (n = 29/30) had the output “coverage too low”, suggesting genome quality limitations (**Figure 13**). The remaining isolate (GPS_KH_COMRU131) was classed as “untypable”, and we confirmed that there was no intact *cps* locus for this isolate via manual investigation. In addition, this isolate was determined to have a non-typable serotype by latex agglutination.

Overall, we only identified 84 (0.5%; n = 84/17,933) isolates within the dataset that have phenotypically determined serotypes that do not match SeroBA(v2.0) or PneumoCaT results at the serotype or serogroup level.

We manually investigated the *cps* region for each of these 84 isolates to determine which serotyping method was correct (**Supplementary Data 6**). For 91.7% of these mismatches (n = 77/84), we found an intact *cps* region for the serotype predicted by SeroBA(v2.0), but not for the phenotypically determined serotype. Therefore, SeroBA(v2.0) was determined to be correct in these cases. These mismatches could have occurred due to mixed samples containing multiple pneumococcal serotypes, the mislabelling or mishandling of tubes during DNA extraction/serotyping, or the subjectivity of Quellung/Latex agglutination results [43]. In 6.0% (n = 5/84) of the mismatches, a read quality issue prevented SeroBA from making a serotype prediction. In only 2.3% (n = 2/84) of the mismatches (0.01% of the isolates with phenotypic serotype data; n = 2/17,933) SeroBA was incorrect, as we found an intact *cps* region for the phenotypically determined serotype but not for the serotype predicted by SeroBA(v2.0) (GPS_ZA_CARRIAGE_SP922, GPS_BR_1063_12_R1). This shows that overall SeroBA(v2.0) has a very high concordance with phenotypic serotypes, and so is an accurate bioinformatics serotyping tool.

To ensure reproducibility, SeroBA(v2.0) was run on the 26,306 genomes three times and the *in silico* serotype and genetic subtypes outputs were compared between the runs. The outputs were found to be 100% identical.

## Discussion

As WGS has become more affordable and accessible, there has been a wealth of genomic data generated; the best example of this is the GPS project which has WGS and phenotypic data for over 26,000 pneumococcal isolates [42]. This has led to a shift in serotyping methodology towards using WGS data to report pneumococcal serotypes based on only genetic information. SeroBA(v2.0) provides a means to take WGS data and understand how the genetic information relates to the phenotypic properties of each serotype i.e. how the body sees each serotype. This is particularly important for understanding the properties of novel serotypes; in just the past four years, nine new serotypes have been discovered. Accurate and timely serotyping of pneumococcal isolates is imperative to monitor the variation within the global pneumococcal population. In this work, we present a free online resource, the SeroBAnk, a knowledge hub of information about all known pneumococcal serotypes (https://www.pneumogen.net/gps/#/serobank). For each serotype, a reference sequence, visualisation of the *cps* locus, capsule structure, and information about which vaccines include the serotype are available. This resource aims to provide an up to date repertoire of pneumococcal serotypes, serving as a foundation for further discoveries in capsular antigens of pneumococci. It will facilitate the global pneumococcal research community in building upon existing knowledge and contributing to the shared knowledge hub.

From this, we developed SeroBA(v2.0) as an updated version of SeroBA [23]. As current bioinformatics serotyping tools are infrequently updated, none can type the most recently discovered serotypes. However, SeroBA(v2.0) can identify 102 of the 107 serotypes described to date, as well as 18 genetic subtypes. 5 serotypes cannot be identified due to either having an inconsistent genetic basis or no reference sequence being available. This provides the community with an up to date tool to reliably serotype pneumococcal isolates from whole genome sequencing data. Genetically-predicted serotyping data from SeroBA(v2.0) should be used in tandem with phenotypically-determined serotyping data. There are limitations to all WGS based serotyping tools (including SeroBA); for example, they cannot assess the functionality of the genes of the *cps* locus due to the high computational cost of assembling all the genes in the *cps* locus.

We propose that to keep SeroBA(v2.0) updated, members of the pneumococcal community are encouraged to submit novel serotypes as they are discovered. These can then be incorporated into the SeroBAnk and SeroBA(v2.0) with relative ease, ensuring that they remain up to date and relevant tools for studying *S. pneumoniae*. New serotypes can be submitted for addition to SeroBA(v2.0) via the issues page to the SeroBA(v2.0) GitHub repository (https://github.com/GlobalPneumoSeq/seroba/issues). Manual curation will be conducted before addition to the SeroBAnk. A novel serotype should demonstrate a distinct *cps* locus, capsular structure, and serological profile, and will rely on collaboration between different lab groups [10].

We have created the SeroBAnk resource, which highlights the genetic basis for each serotype, and updated SeroBA(v2.0) to be able to accurately serotype 102 of the 107 currently described pneumococcal serotypes. Being able to reliably serotype pneumococcal isolates from whole genome sequencing data is a key part of tracking and understanding the evolution and pathogenicity of *S. pneumoniae,* as well as for monitoring the changes to serotype distributions following vaccine introduction. Therefore, the long-term maintenance of both resources is key. We therefore propose a community-centred update system to ensure that when new serotypes are discovered, they are added to the SeroBAnk and SeroBA(v2.0) in a timely manner. This will ensure that new serotypes can always be genetically predicted from whole genome sequencing data, which is crucial for understanding the distribution of serotypes within a population.

## Conflicts of Interest

The authors declare that there are no conflicts of interest. All co-authors have seen and agree with the contents of the manuscript and there is no financial interest to report. The findings and conclusions in this manuscript are those of the authors and do not necessarily represent the official position of the US Centers for Disease Control and Prevention.

## Funding Information

This work was supported by Wellcome under grant reference 206194 and by the Bill & Melinda Gates Foundation under Investment ID INV-003570. CS was supported by a Rebecca Cooper Fellowship from the Rebecca L Cooper Medical Research Foundation. Authors affiliated with the Murdoch Children’s Research Institute were supported by the Victorian Government’s Operational Infrastructure Support Program. The funding sources had no role in analysis or data interpretation. For the purposes of Open Access, the author has applied a CC BY public copyright licence to any Author Accept Manuscript version arising from this submission. The corresponding author had full access to the data and is responsible for the final decision to submit.

## Acknowledgements

We would like to thank all members of The Global Pneumococcal Sequencing Consortium for their collaborative spirit and determination during the task of sampling, extracting, and sequencing this dataset. Finally, we would like to thank the Pathogen Informatics Team at the Wellcome Sanger Institute for technical support of bioinformatics analyses.

## Notes

https://github.com/GlobalPneumoSeq/seroba

### Competing Interest Statement

The authors have declared no competing interest.

