## Supplementary1 for "SeroBA(v2.0) and SeroBAnk: a robust genome-based serotyping scheme and comprehensive atlas of capsular diversity in Streptococcus pneumoniae"

### Introduction

This document contains summary tables describing the genetic basis for determining the new serotypes and subtypes added to the SeroBA CTVdb. It is intended to be used in addition to the original PneumoCAT reference determination document which was last updated in 2019. SeroBA still uses all the genetic determinants from this document for determining the serotypes described in the PneumoCAT reference determination document.

This information is also available in the yaml files in the SeroBA GitHub repository (https://github.com/sanger-bentley-group/seroba)

This document contains summary tables describing the genetic and structural differences for the new serotypes and subtypes added to the CTVdb.

Feedback is welcome, you can get in touch via the GitHub repository and you can customise the CTVdb to suit your own needs.

#### New Serotype Summary Table

| Serogroup | Serotype | Distinguishing genetic features | Functional effect |
|---|---|---|---|
| 11 | 11A/11B/11C/11D /11E/11F/11F_like | Genes *wcwC* and *wcjE* are present in 11A, 11D, 11E and 11F_like, whereas *wcwR* is present in 11B and 11C. | *wcwC*, *wcjE* and *wcwR* are acetyl transferases – differences in acetylation |
| | 11A/11B/11C/11D /11E/11F/11F_like | Frameshift mutation delA 130 in *gct* in 11B and 11F causes inactivation of *gct* gene | Presence of Gro-1P correlates with intact *gct* gene (Mavrodi et al. 2007) |
| | 11A/11D/11E/11F /11F_like | SNPs in *wcrL* (Table 1) | Donor sugar for *wcrL* is GlcpNAc in types 11F, 11B and 11C, Glcp in 11A (Mavrodi et al. 2007) |
| | 11F_like | Divergent *wcrL* and *wcwC* genes | *wcwC* changes not understood yet, lower glycosyltransferase activity in *wcrL* compared to 11F (Manna et al. 2018) |
| 15 | 15A/15B/15C/15 D/15F | 15F has 4 additional genes: *glf*, *rmlB*, *rmlD* and *wcjE* | *glf*, *rmlB* and *rmlD* are involved in sugar biosynthesis. *wcjE* is an acetyltransferase |
| | 15A/15B/15C/15 D | 15A/15D *wzd* has ~70% identity in the last 300bps compared to *wzd* 15B/15C | *wzd* is involved in translocation of the CPS to the cell surface, thus determining the length of the capsule polysaccharide chain |
| | 15B/15C | Difference in TA tandem repeat near position 413 in *wciZ*, leading to frameshift in 15C (Bentley et al. 2006) | Differences in acetylation 15B, 15C and 15B/C results can be assigned |
| | 15A/15D | Difference in *wzy* allele – 15A *wzy-15A*. 15D – *wzy-15D* (a *wzy-15AF* hybrid) | Different polymerization linkages (Mavrodi et al. 2007) |

| | | | |
|---|---|---|---|
| 20 | 20A/20B | Insertion in polyA tract in *whaF* near position 881 leads to truncated gene in 20A | Loss of glycosyltransferase function in 20A (Calix et al. 2012) |
| | 20C | Disruptive mutation in *wciG* leads to truncated gene | Loss of O-acetyl transferase function in 20C (unpublished) |
| 24 | 24A / 24B/24C/24F | 24A lacks *rbsF* gene present in 24B/24C/24F | Loss of ribofuranose biosynthesis in 24A (Ganaie et al. 2021) |
| 33 and 37 | 33A/33E/33F/37 | 37 carries *tts* (a transferase gene) | *tts* is responsible for polysaccharide capsule biosynthesis (Waite et al. 2003) |
| | 33A/33F | Frameshift mutation insT 433 in *wcjE* gene | Loss of acetyltransferase function leads to differences in acetylation (Mavrodi et al. 2007) |
| | 33B/33D | *wciN*$\alpha$ in 33B, *wciN*$\beta$ in 33D | *wciN*$\alpha$ codes for a glycosyltransferase, *wciN*$\beta$ codes for a galactosyltransferase |
| | 33A/33E/33F | Nonsense mutation in *wciE*, around position 273 C->T truncates *wciE* in 33E | Inactivated glycosyltransferase gene in 33E (Ganaie et al. 2023) |
| 33G/33H | 33G/33H | 33H has a disruptive mutation in *wciG* causing inactivation | Loss of O-acetyl transferase function in 33H (unpublished) |
| 36 | 36A/36B | *wcjA*$\alpha$ in 36A, *wcjA*$\beta$ in 36B. SNPs in *wcjA* (Table 6) | Mutations lead to the incorporation of Glcp in 36A vs Galp in 36B (Ganaie et al. 2023) |

#### New subtype summary table

| Serotype | Subtypes | Distinguishing genetic features | Functional effect |
|---|---|---|---|
| 6A | 6A/6A-I/6A-II/6A-III/6A-IV/6A-V/6A-VI | Except for 6A-IV, different allelic combinations define each subtype within the 6A serotype | Nonsynonymous mutations in all subtypes leading to varying levels of structural divergence from the 6A reference sequence |
| | 6A-I | 3 divergent alleles: *wzg*$\alpha$, *rmlA-3* and *rmlB-4* | |
| | 6A-II | 3 divergent alleles: *wzg*$\beta$, *wzy*$\alpha$ and *rmlA-2* | |
| | 6A-III | 2 divergent alleles: *wzg*$\chi$ and *rmlA-4* | |
| | 6A-IV | SNP in *wze* pos 488 | |
| | 6A-V | 2 divergent alleles: *wzg*$\alpha$ and *rmlA-5* | |
| | 6A-VI | 4 divergent alleles: *wzg*$\beta$, *wzy*$\alpha$, *rmlA-2* and *rmlC-2* | |
| 6B | 6B/6B-I/6B-II | Different allelic combinations define each subtype within the 6B serotype | Nonsynonymous mutations in all subtypes leading to varying levels of structural divergence from the 6B reference sequence |
| | 6B-I | 3 divergent alleles: *wzg*$\chi$, *rmlA-4* and *rmlB-3* | |
| | 6B-II | 3 divergent alleles: *wzg*$\delta$, *rmlA-5* and *rmlB-3* | |
| 19A | 19A/19A-I/19A-II/19A-III/19AF | Different allelic and mutational combinations define each subtype within the 19A serotype | Nonsynonymous mutations in all subtypes leading to varying levels of structural divergence from the 19A reference sequence |
| | 19A-I | 2 divergent rhamnose alleles: *rmlB-2* and *rmlD-2* | |

| | | | |
|---|---|---|---|
| | 19A-II | 2 divergent rhamnose alleles: *rmlB-2* and *rmlD-2*. SNP in *wzg* differentiates from 19A-I | |
| | 19A-III | SNP in *wzd* pos 154 determines 19A-III | |
| | 19AF | 19AF has 19F *wzy* allele | 19AF phenotype as 19F despite having overall 19A-like capsular operon sequence |
| 19F | 19F/19F-I/19F-II/19F-III/19F-IV | Different allelic and mutational combinations define each 19F subtype | Nonsynonymous mutations in all subtypes leading to varying levels of structural divergence from the 19F reference sequence |
| | 19F-I | SNPs in *wze* | |
| | 19F-II | SNPs in *wze* and divergent wchA-4 allele | |
| | 19F-III | SNPs in *wchO*, *wzx* and *wzy* | |
| | 19F-IV | 3 divergent alleles: *wchA-4, rmlB-6* and *wzg-2* | |

Sequences for the above table were obtained from Elberse et al. 2011. See Table 14.

#### Serogroup 11

11F_like was previously typed as 11A by SeroBA

***Table 1***

| Gene | variant | 11A | 11B | 11C | 11D | 11E | 11F | 11F_like |
|---|---|---|---|---|---|---|---|---|
| ***wcwC*** | detected | Y | N | N | Y | Y | Y | Y |
| ***wcjE*** | detected | Y | N | N | Y | Y | Y | Y |
| ***wcwR*** | detected | N | Y | Y | N | N | N | N |
| ***gct*** | pseudo (130delA) | N | Y | N | N | N | Y | N |
| ***wcrL*** | pos 49 | TCA | - | - | TCA | TCA | TCA | CCA |
| ***wcrL*** | pos 334 | AAT | - | - | ACT | AAT | GCT | AAT |
| ***wcrL*** | Pos 493 | GAC | - | - | GAC | GAC | GAC | AAT |

#### Serogroup 15

15D was previously typed as 15F or 15A by SeroBA

***Table 2***

| Gene | variant | 15A | 15B | 15C | 15D | 15F |
|---|---|---|---|---|---|---|
| ***wzd*** | allele | wzdα | wzdβ | wzdβ | wzdα | - |
| ***glf*** | detected | N | N | N | N | Y |
| ***rmlB*** | detected | N | N | N | N | Y |
| ***rmlD*** | detected | N | N | N | N | Y |
| ***wcjE*** | detected | N | N | N | N | Y |
| ***wciZ*** | pseudo | - | N, [412,417] | Y, [412,417] | - | - |
| ***wzy*** | allele | wzyα | - | - | wzyβ | - |

#### Serogroup 20

Serogroup 20 isolates were typed at the serogroup level only in the old version of SeroBA.

***Table 3***

| Gene | variant | 20A | 20B | 20C |
|---|---|---|---|---|
| ***whaF*** | pseudo | Y | N, [881,882] | N, [881,882] |
| ***whaF*** | pseudo | N | N | N |
| ***wciG*** | pseudo | N | N | Disruptive mutation |

#### Serogroup 24

Serogroup 24 isolates were previously typed as “serogroup 24”. 24A can now be distinguished due to the absence of the rbsF gene. The genetic determination of 24B/C/F was deemed too difficult due to a lack of knowledge about the specific genetic determinants (mutations) for these serotypes.

***Table 4***

| Gene | variant | 24A | 24B/24C/24F |
|---|---|---|---|
| ***rbsF*** | detected | Y | N |

#### Genogroup 33A/33E/33F/37

33E was previously typed as 33F in SeroBA.

***Table 5***

| Gene | variant | 33A | 33E | 33F | 37 |
|---|---|---|---|---|---|
| ***tts*** | detected | N | N | N | Y |
| ***wcjE*** | pseudo | N, [[433, 434], T]] | - | Y, [[433, 434], TA]] | - |
| ***wciE*** | pseudo | N, [[273, 274, C]] | Y, [[273, 274, T]] | N, [[273, 274, C]] | - |

#### Genogroup 33G/33H

***Table 6***

| Gene | variant | 33G | 33H |
|---|---|---|---|
| ***wciG*** | pseudo | N | Disruptive mutation |

#### Serogroup 36

In the previous version of SeroBA, serogroup 36 samples were serotyped at the serogroup level only.

***Table 7***

| Gene | variant | 36A | 36B |
|---|---|---|---|
| ***wcjA*** | allele | wcjA$\alpha$ | wcjA$\beta$ |
| ***wcjA*** | pos 430 | [AGT, S] | [GGT, G] |
| ***wcjA*** | pos 805 | [TTT, F] | [ATG, M] |
| ***wcjA*** | pos 808 | [GGC, G] | [GCT, A] |
| ***wcjA*** | pos 817 | [GCT, A] | [GTT, V] |

#### Serotype 6A

Samples which are typed as 6A now undergo subtyping to highlight the genetic diversity within the 6A serotype. Samples are only subtyped if all allelic and mutational criteria are satisfied.

***Table 8***

| Gene | variant | 6A | 6A-I | 6A-II | 6A-III | 6A-IV | 6A-V | 6A-VI |
|---|---|---|---|---|---|---|---|---|
| ***wzg*** | allele | wzg$\gamma$ | wzg$\alpha$ | wzg$\beta$ | wzg$\chi$ | wzg$\gamma$ | wzg$\alpha$ | wzg$\beta$ |
| ***rmlA*** | allele | rmlA-1 | rmlA-3 | rmlA-2 | rmlA-4 | rmlA-1 | rmlA-2 | rmlA-4 |
| ***rmlB*** | allele | rmlB-1 | rmlB-4 | rmlB-1 | rmlB-1 | rmlB-1 | rmlB-1 | rmlB-1 |
| ***rmlC*** | allele | rmlC-1 | rmlC-1 | rmlC-1 | rmlC-1 | rmlC-1 | rmlC-1 | rmlC-2 |
| ***wzy*** | allele | wzy$\gamma$ | wzy$\gamma$ | wzy$\alpha$ | wzy$\gamma$ | wzy$\gamma$ | wzy$\gamma$ | wzy$\alpha$ |
| ***wze*** | pos 488 | [GCT, A] | [GCT, A] | [GCT, A] | [GCT, A] | [GTT, V] | [GCT, A] | [GCT, A] |

#### Serotype 6B

Samples which are typed as 6B now undergo subtyping to highlight the genetic diversity within the 6B serotype. Samples are only subtyped if all allelic and mutational criteria are satisfied.

*Table 9*

| Gene | variant | 6B | 6B-I | 6B-II |
|---|---|---|---|---|
| ***wzg*** | allele | wzg$\varepsilon$ | wzg$\chi$ | wzg$\delta$ |
| ***rmlA*** | allele | rmlA-3 | rmlA-4 | rmlA-5 |
| ***rmlB*** | allele | rmlB-4 | rmlB-3 | rmlB-3 |

#### Serotype 19A

Samples which are typed as 19A now undergo subtyping to highlight the genetic diversity within the 19A serotype. Samples are only subtyped if all allelic and mutational criteria are satisfied. In some cases, it is not possible to determine the SNPs which differentiate 19A-I from 19A-II, when this occurs the type called is 19A-I/19A-II.

***Table 10***

| Gene | variant | 19A | 19A-I | 19A-II | 19A-III | 19AF |
|---|---|---|---|---|---|---|
| ***rmlB*** | allele | rmlB-1 | rmlB-2 | rmlB-2 | rmlB-1 | rmlB-1 |
| ***rmlD*** | allele | rmlD-1 | rmlD-2 | rmlD-2 | rmlD-1 | rmlD-1 |
| ***wzy*** | allele | wzy-1 | wzy-1 | wzy-1 | wzy-1 | wzy-2 |
| ***wzg*** | pos 445 | [GCT, A] | [TCT, S] | [GCT, A] | [GCT, A] | [GCT, A] |
| ***wzg*** | pos 484 | [AAT, N] | [AAT, N] | [GAC, D] | [AAT, N] | [AAT, N] |
| ***wzd*** | pos 154 | [ACC, T] | [ACC, T] | [ACC, T] | [ATC, I] | [ACC, T] |

#### Serotype 19F

Samples which are typed as 19F now undergo subtyping to highlight genetic diversity. Samples are only subtyped If all allelic and mutational criteria are satisfied.

***Table 11***

| Gene | variant | 19F | 19F-I | 19F-II | 19F-III | 19F-IV |
|---|---|---|---|---|---|---|
| ***rmlB*** | allele | rmlB-5 | rmlB-5 | rmlB-5 | rmlB-5 | rmlB-6 |
| ***wchA*** | allele | wchA-3 | wchA-3 | wchA-4 | wchA-3 | wchA-4 |
| ***wzg*** | allele | wzg-1 | wzg-1 | wzg-1 | wzg-1 | wzg-2 |
| ***wchO*** | pos 67 | [AGT, S] | [ATT, I] | [AGT, S] | [ATT, I] | [AGT, S] |
| ***wze*** | pos 135 | [GAG, E] | [AGG, R] | [GGG, G] | [GAG, E] | [GGG, G] |
| ***wze*** | pos 207 | [ATC, I] | [ATC, I] | [ATC, I] | [ATC, I] | [CTC, L] |
| ***wze*** | pos 477 | [ATT, I] | [GTT, V] | [ATT, I] | [ATT, I] | [ATT, I] |
| ***wzx*** | pos 1111 | [ATA, I] | [ATG, M] | [ATA, I] | [ATG, M] | [ATA, I] |
| ***wzx*** | pos 1133 | [ATA, I] | [CTA, L] | [CTA, L] | [CTA, L] | [CTA, L] |
| ***wzy*** | pos 213 | [ATT, I] | [CTT, L] | [ATT, I] | [CTT, L] | [ATT, I] |

#### Serotype 33F

***Table 12***

| Gene | variant | 33F | 33F-1a | 33F-1b |
|---|---|---|---|---|
| ***wcjE*** | detected | Y | N | N |
| ***wchO*** | pseudo | NA | N | Y |

#### Serotype Reference sequence information

***Table 13***

| Serotype | Accession number | Strain | Reference |
|---|---|---|---|
| **11F_like** | MF140334 | PMP1342 | Manna et al. 2018 |
| **15D** | SAMN14150919 | 20184245 | Pimenta et al. 2021 |
| **20A** | JQ653094 | 6320 | Calix et al. 2012 |
| **20B** | JQ653093 | 5931-06 | Calix et al. 2012 |
| **20C** | ERS653863 | GPS_NP_6772 | Unpublished |
| **24C** | MW683289 | MNY576 | Ganaie et al. 2021 |
| **33E** | SAMEA2203953 | GPSC3 | Ganaie et al. 2023 |
| **33H** | ERS464499 | GPS_ZA_887 | Unpublished |
| **36A** | CR931708 | 1095/39 | Bentley et al. 2006 |
| **36B** | MK606436 | GPS_NP_1196 | Ganaie et al. 2023 |

#### Subtype reference sequence information

***Table 14***

| Subtype | Accession number | Reference |
|---|---|---|
| **6A-I** | JF911487 | Elberse et al. 2011 |
| **6A-II** | JF911488 | Elberse et al. 2011 |
| **6A-III** | JF911497 | Elberse et al. 2011 |
| **6A-IV** | JF911492 | Elberse et al. 2011 |
| **6A-V** | JF911491 | Elberse et al. 2011 |
| **6A-VI** | JF911490 | Elberse et al. 2011 |
| **6B-I** | JF911498 | Elberse et al. 2011 |
| **6B-II** | JF911499 | Elberse et al. 2011 |
| **19A-I** | JF911512 | Elberse et al. 2011 |
| **19A-II** | JF911514 | Elberse et al. 2011 |
| **19A-III** | JF911520 | Elberse et al. 2011 |
| **19F-I** | JF911525 | Elberse et al. 2011 |
| **19F-II** | JF911524 | Elberse et al. 2011 |
| **19F-III** | JF911522 | Elberse et al. 2011 |
| **19F-IV** | JF911528 | Elberse et al. 2011 |

| | | |
|---|---|---|
| **33F-1a** | GPS_US_PATH361 8 | Ganaie et al. 2023 |
| **33F-1b** | MK606435 | Ganaie et al. 2023 |

#### Non encapsulated reference strains

NCC1_pspK_non_encapsulated- JF489996
NCC3_aliD_non_encapsulated - JF490008
NCC2_aliC_aliD_non_encapsulated - HE651292
NCC2_S_mitis_aliC_aliD_non_encapsulated - HE651274
